# Parvalbumin interneurons mediate spontaneous hemodynamic fluctuations

**DOI:** 10.1101/2025.08.15.670614

**Authors:** Adiya Rakymzhan, Mitsuhiro Fukuda, Alberto L. Vazquez

## Abstract

Resting-state hemodynamic fluctuations are closely linked to gamma-band neural activity, yet the cellular drivers of this neurovascular coupling remain unclear. Given their established contribution in generating gamma oscillations, parvalbumin (PV) interneurons are prime candidates for regulating spontaneous cerebral blood flow (CBF) dynamics. Using chemogenetic tools in awake PV-Cre mice, we modulated PV interneuron activity and measured effects on neural network activity, local field potentials (LFP), and hemodynamics. Two-photon (2P) calcium imaging confirmed effective PV modulation, which affected excitatory neuron activity. PV suppression reduced high-gamma power, increased low-frequency LFP activity, and elevated basal CBF. 2P vessel imaging showed increased basal arterial diameter and significantly greater diameter fluctuation power in deeper cortical layers enriched with PV cells, but not in superficial layers. PV suppression also significantly weakened the correlation between gamma LFP power and CBF. Despite increased apparent neuronal synchrony during PV suppression, its relationship to arterial dynamics remained stable, possibly due to compensatory regulation by subsets of PV-positive and PV-negative cells. These findings provide causal evidence that PV interneuron contribute to spontaneous neurovascular dynamics and mediate the link between gamma oscillations and resting-state hemodynamic signals, revealing their significant role in maintaining functional connectivity and vascular regulation during non-task-engaged brain states.

## INTRODUCTION

Spontaneous fluctuations in hemodynamic signals, particularly at frequencies centered around 0.1 Hz, appear during so-called resting state [1–4]. This term typically refers to a condition where a subject lies in the functional magnetic resonance imaging (fMRI) scanner without performing an explicit task or being exposed to a specific stimulus. Human fMRI studies and optical imaging studies in animals revealed that these low-frequency fluctuations are strongly synchronized within like networks and across distant connected regions [1–8]. This phenomenon, known as ‘‘functional connectivity’’, offers valuable insights into the brain’s functional organization, making hemodynamic-based imaging techniques an important tool for studying brain function [2, 6, 9–10].

Hemodynamic synchronized activity is generally associated with neural activity in known anatomical connections. Simultaneous fMRI and intracortical electrophysiological recordings in the monkey visual cortex during visual stimulation have shown that local field potentials (LFPs) provide a better estimate of blood-oxygen-level-dependent (BOLD) responses than multi-unit activity [11]. To compare neural activity with slower hemodynamic fluctuations that typically lag behind neural activity by a few seconds, researchers have introduced band-limited power (BLP) signals derived from LFPs [12]. These signals exhibit large-amplitude fluctuations over extended time scales and are also synchronized across distant cortical regions [12]. During resting state, concurrent fMRI-LFP recordings in monkeys have demonstrated that spontaneous BOLD fluctuations strongly correlate with BLP power in the gamma frequency range [13–14]. Similarly, in mice, low-frequency arterial oscillations have been found to phase-lock with slow variations in gamma BLP [4, 8, 15]. These studies highlight a robust relationship between neural gamma activity and spontaneous hemodynamic fluctuations. However, the mechanisms driving this correlation remain poorly understood.

Gamma activity in the cortex is thought to have strong contribution from parvalbumin (PV) interneuron activity. These interneurons provide strong inhibitory input to local populations of asynchronously firing principal cells, effectively suppressing their activity and generating synchronized gamma rhythm within the network [16]. Optogenetic studies have further demonstrated the critical role of PV interneuron in gamma oscillation generation, where selective activation of PV cells induced gamma rhythms and their inhibition attenuated them significantly [17–18]. Given the strong correlation between gamma BLP and spontaneous hemodynamic fluctuations, it is plausible that PV interneuron not only drive gamma activity but also play a key role in regulating resting-state cerebral blood flow (CBF) dynamics.

Neurons regulate CBF by manipulating the tone of nearby blood vessels, a process known as neurovascular coupling (NVC). Different neuronal populations exert distinct effects on vascular dynamics. Advances in genetic manipulation tools and the emergence of optogenetics in rodent models have enabled researchers to dissect the contributions of specific neuronal subpopulations to CBF regulation. In vivo optogenetic studies in anesthetized mice have shown that activating GABAergic inhibitory neurons induces a larger increase in CBF than activating excitatory neurons [19]. Among inhibitory neurons, our previous work in awake mice demonstrated that activating PV interneuron produces slow, gradual vasodilations, particularly in deeper cortical layers where these neurons are predominantly concentrated [20–21]. Notably, these slow vasodilations closely resemble low-frequency, high-amplitude hemodynamic fluctuations observed in rodents and humans during the resting state. While studies on evoked activity of PV neurons have provided valuable insights into their influence on CBF, their role in regulating spontaneous CBF fluctuations during awake resting-state activity remains unclear [20–25]. Correlational analyses suggest that synchronized PV cell activity drives depth-specific vascular changes, but more direct causal evidence is lacking [21].

In this study, we sought to provide more direct evidence for the role of PV interneuron in NVC during resting state by selectively modulating their activity using chemogenetics. To achieve this, we used transgenic PV-cre mice to express cre-dependent Designer Receptors Exclusively Activated by Designer Drugs (DREADD), enabling selective modulation of PV interneurons in the primary somatosensory cortex (S1). Our first objective was to assess the contribution of PV interneurons to network dynamics by imaging calcium activity in PV-positive (PV+) and PV-negative (PV-) cells using 2P and measuring LFP activity using electroencephalogram (EEG) at the baseline and after DREADD activation. The second goal was to evaluate how PV cell modulation affects spontaneous hemodynamic fluctuations by measuring CBF changes with Laser Doppler flowmetry (LDF) and vessel tone with 2P. Finally, we investigated the impact of PV suppression on the coupling between neural activity and hemodynamic oscillations during resting state. Our findings indicate that PV interneuronssignificantly modulate both ongoing network activity, spontaneous hemodynamic fluctuations, and NVC. These results suggest that PV cell signaling significantly contributes to the mechanism underlying the strong correlation between gamma oscillations and CBF dynamics during resting-state conditions.

## Materials and Methods

### Experimental model

We used a total of 17 adult transgenic mice (male n = 9, female n = 8) expressing recombinase in PV neurons (strain B6.129P2-Pvalb(tm1-cre)Arbr/J, breeders were purchased from The Jackson Laboratory; Bar Harbor, ME USA). All procedures performed followed an experimental protocol approved by the University of Pittsburgh Institutional Animal Care and Use Committee (IACUC) by the standards for humane animal care and use as set by the Animal Welfare Act (AWA), the National Institute of Health Guide for the Care and Use of Laboratory Animals and ARRIVE (Animal Research: Reporting in Vivo Experiments).

### Animal surgery for awake head-fixed experiments

When mice were 8-10 weeks of age, we injected cre-dependent AAV encoding inhibitory DREADD-mCherry (AAV-hSyn-DIO-hM4D(Gi)-mCherry) or excitatory DREADD-mCherry (AAV-hSyn-DIO-hM3D(Gq)-mCherry; University of North Carolina Vector Core, Chapel Hill, NC USA) in S1 (Figure 1A-C). To drive the expression of calcium (Ca^2+^) reporter in neurons using the Synapsin promoter, mice were injected with AAV-Syn-GCaMP7f or AAV-Syn-GCaMP8f (Addgene, Watertown, MA USA) in the same region of S1 (Figure 1A-C). For the control group, we used 3 male adult PV-cre mice injected with AAV-hSyn-DIO-mCherry and AAV-Syn-GCaMP7f or AAV-Syn-GCaMP8f.

**Figure 1.**
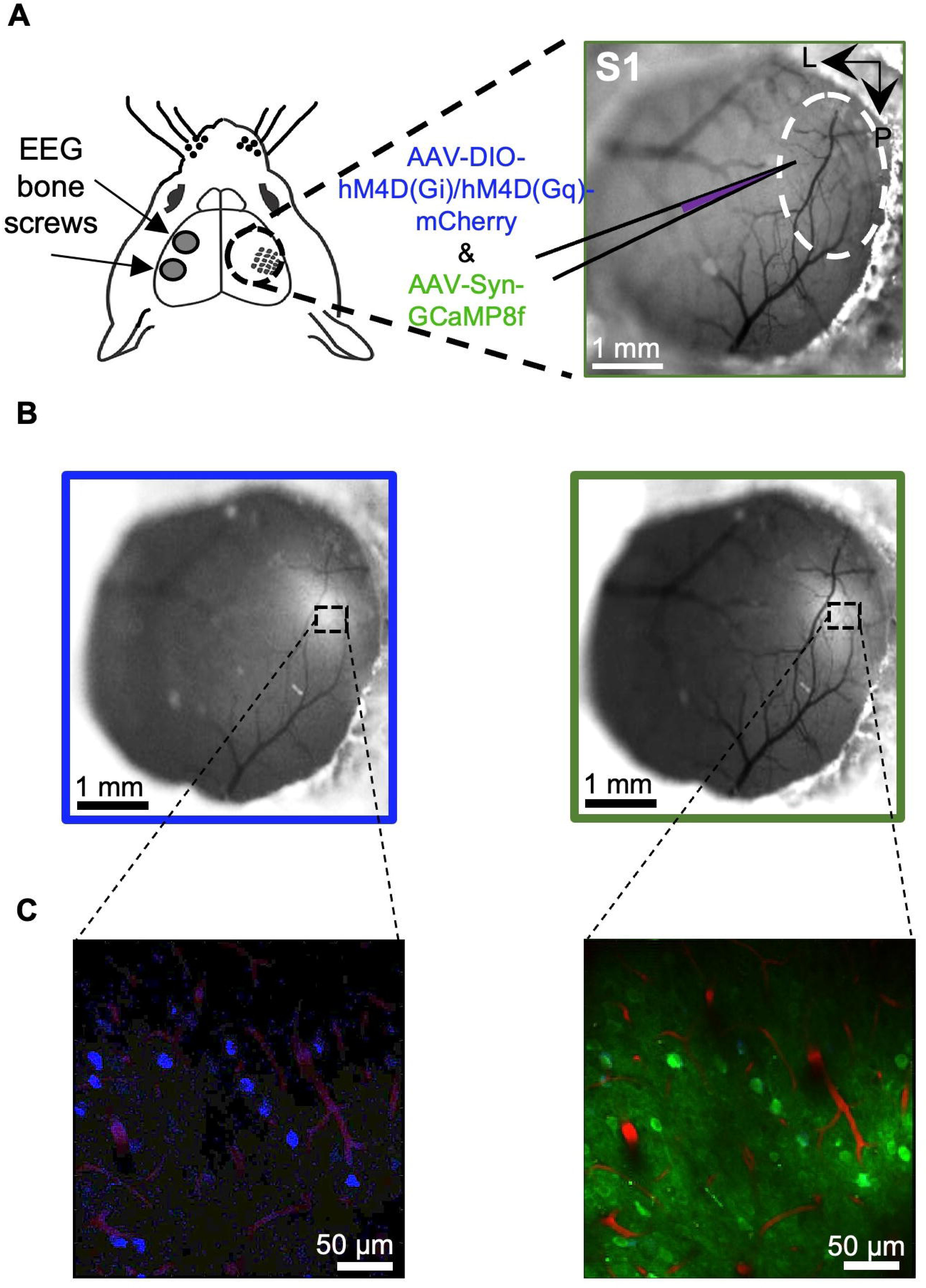
Animal Preparation. **A**. Cranial window implanted in S1 of a PV-cre mouse, where we injected inhibitory DREADD (AAV-DIO-hM4D(Gi)-mCherry) and calcium reporter (AAV-Syn-GCaMP8f) viruses. Two metal EEG bone screws were implanted in the opposite hemisphere for EEG recordings. **B.** Surface fluorescence microscope images below showing the regions expressing hM4D(Gi)-mCherry (blue box) and GCaMP8f (green box). **C**. 2P microscopy images of S1 intra-cortical sections (∼250 um depth) showing the regions expressing hM4D(Gi)-mCherry (blue cells) and GCaMP8f (green cells). Vessel contrast enhanced by Sulforhodamine 101 injection.

Each animal underwent a surgical procedure to implant a cranial window for optical access (Figure 1A). Briefly, the exposed brain was sealed with a glass coverslip (4 mm), and a custom head-plate was fixed to the skull. Two metal bone screws (stainless steel bone screws, 0.86 mm × 4 mm, Fine Science Tools) were implanted in the opposite hemisphere for EEG recordings (Figure 1A). Mice were given at least 4 weeks to recover post-surgery and before imaging. During the recovery period, mice were acclimated to head fixation on a treadmill (1 hour of treadmill in their cage followed by 1 hour of treadmill under the microscope for 5 days) during the four weeks post-surgery. Mice were imaged at the age of 3-8 months.

### Two-photon Microscopy Imaging

Each mouse underwent at least 3 awake daytime imaging sessions, with 1 week between the sessions. Before imaging, Sulforhodamine 101 (SR101; Thermo Fisher Scientific, Waltham, MA USA), dissolved in saline (5–10 mM), was injected in mice intraperitoneally (IP; 8 μL/g body weight) for imaging the cerebral vasculature [26]. Imaging was performed with InSight X3 widely tunable ultrafast laser (Spectra-Physics, Milpitas, CA USA), coupled to a two-photon fluorescence microscope (2P Plus, Bruker Nano Inc., Billerica, MA USA) with a 16X water immersion objective lens, 0.80 NA (Nikon Inc., Tokyo, Japan) with a maximum field-of-view (FOV) of 1100 × 1100 μm (Figure 2A). Two-photon excitation was performed at 900 nm for imaging GCaMP and SR101 in S1 cortex at depths of 250-450 μm (layers 1-4). The following imaging parameters were used to acquire time series: raster scanning rate at 2.4 μs/pixel; 332 × 332 μm^2^ FOV with 1.66 μm/pixel; 200 × 200 matrix; 5.1 Hz frame rate. We measured arteries and arterioles since traditionally they are considered the major players in blood flow regulation. The selection of the imaged region was based on three parameters, where the FOV includes: penetrating vessels, GCaMP cells, and PV-mCherry cells.

**Figure 2.**
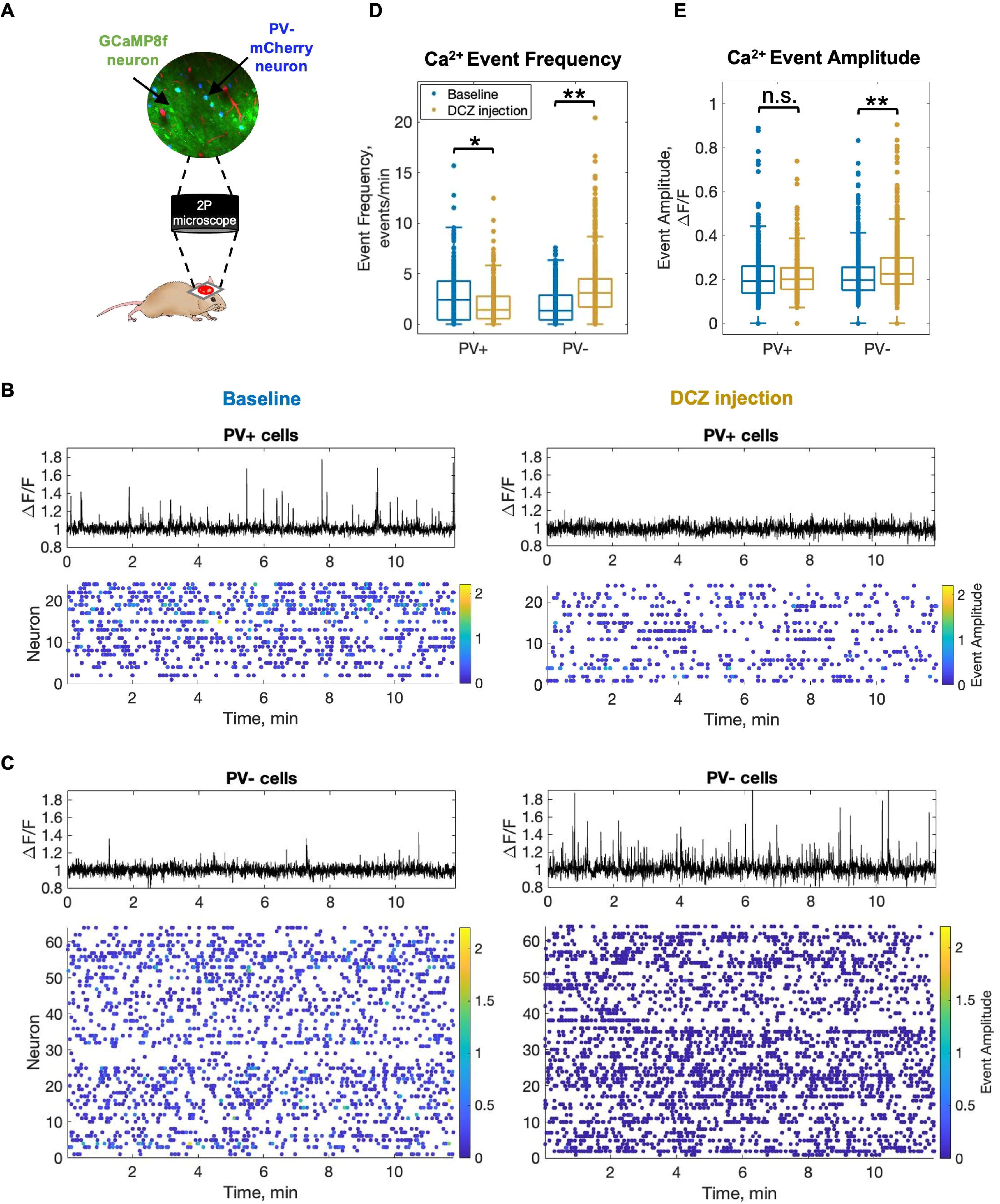
Activation of hM4Di(Gi) DREADD with DCZ effectively suppresses PV neuron activity and enhances network activity. **A.** 2P imaging setup, where neurons expressing GCaMP8f, including PV-mCherry cells, were visualized. **B.** Time profiles of Ca^2+^ fluorescence changes (black traces) in an individual PV+ cell in a PV-hM4Di(Gi) mouse during ongoing activity at baseline and post-DCZ injection. Below are the raster plots of Ca^2+^ activity in PV+ cell population from a single animal at baseline and post-DCZ injection. Each dot represents a Ca^2+^ event and each raw represents an individual neuron. The dot color encodes an event amplitude. **C**. Similar analysis as in B but for PV-cells. **D**. Summary of Ca^2+^ event frequency and **E**. Ca^2+^ event amplitude in PV+ and PV-cells. Significant differences are denoted by * for p<0.05 or ** for p<0.01. Non-significant differences are denoted by n.s.

### EEG and LDF recordings

Electrophysiological activity was recorded with implanted bone screws (Figure 1A, Figure 3A) at a frequency of 1kHz (BIOPAC System, Inc., Goleta, CA). CBF data was acquired with LDF (Periflux 5000/411, Perimed AB, Jarfalla, Sweden) acquired and sampled at 1kHz. The LDF probe, with a tip diameter of 450 μm and operating wavelength of 780 nm, was placed at 30° on the cover glass facing the expressing region, avoiding large surface vessels (Figure 4A).

**Figure 3.**
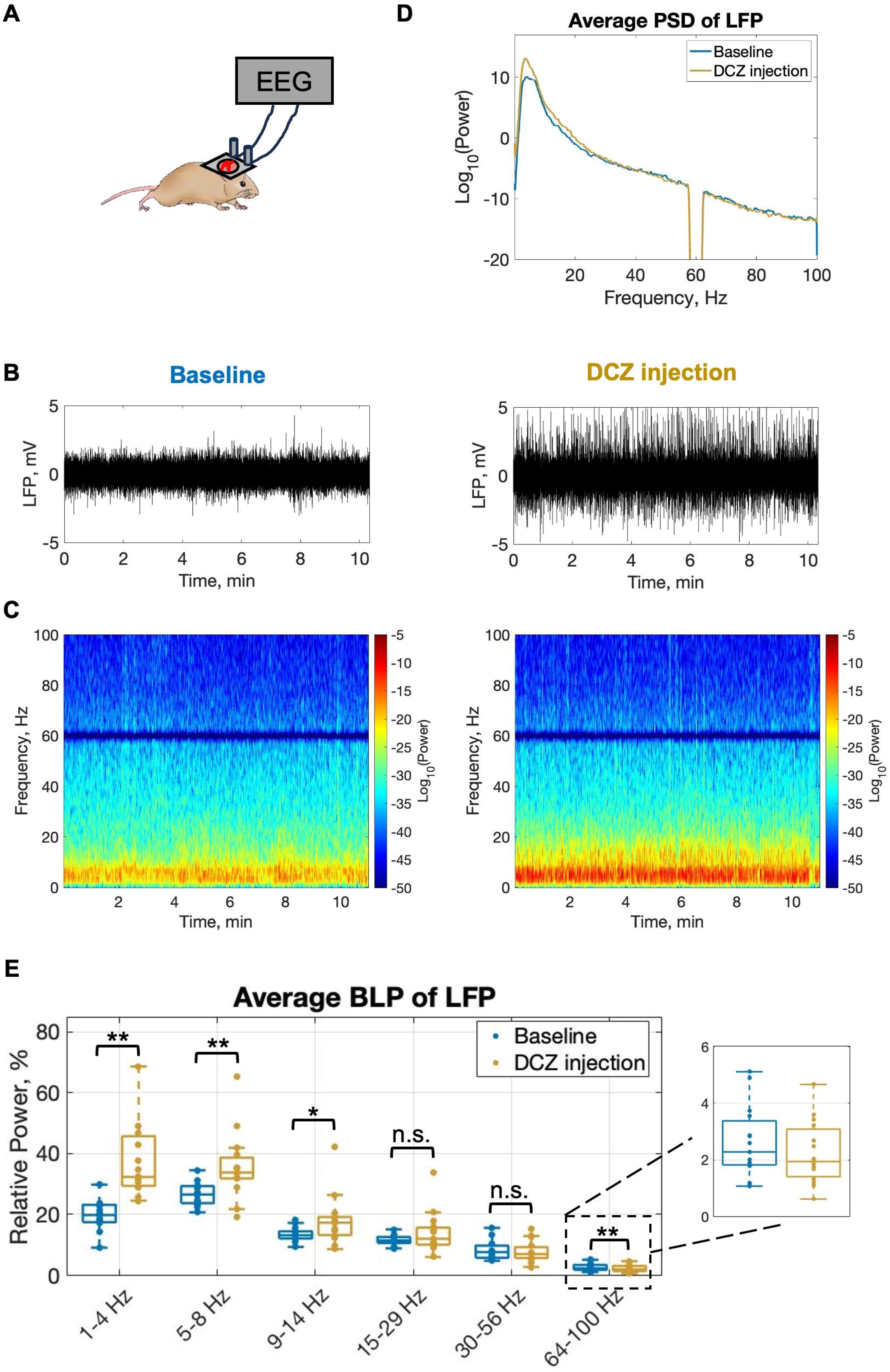
Suppression of PV neuron activity decreases high-gamma power and increases low-frequency power in LFP. **A.** EEG recording setup. **B**. Example of LFP traces and **C**. example of LFP signal spectrogram during ongoing activity at baseline and post-DCZ injection in a PV-hM4Di(Gi) mouse. **D**. Average PSD of LFP signals and **E.** average BLPs of LFP in PV-hM4Di(Gi) mice during ongoing activity at baseline and post-DCZ injection. Significant differences are denoted by * for p<0.05 or ** for p<0.01. Non-significant differences are denoted by n.s.

**Figure 4.**
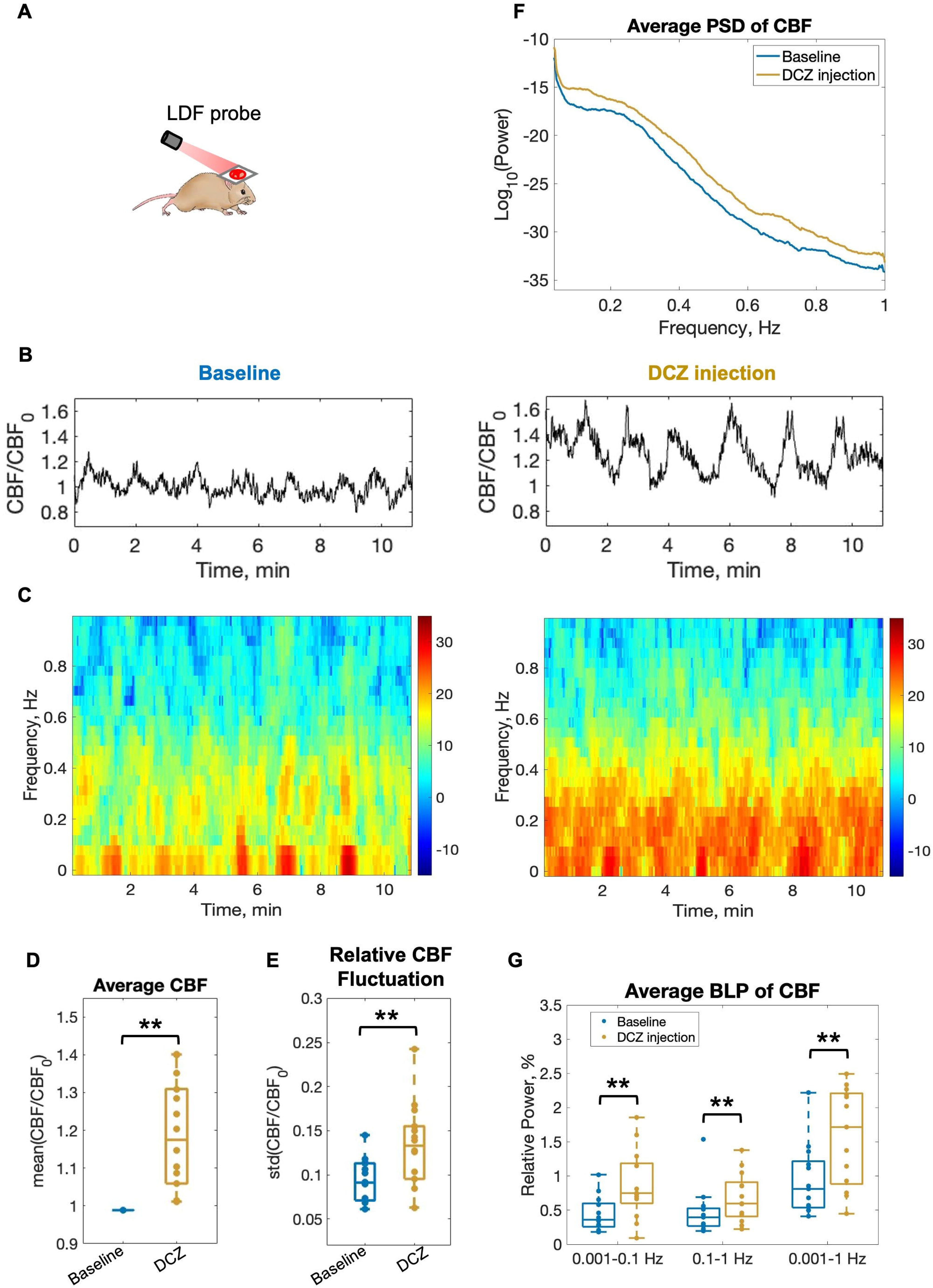
Suppression of PV neuron activity increases spontaneous CBF. **A.** LDF recording setup for CBF measurements, where LDF probe was placed on top of cranial window. **B.** Example of CBF time profiles and **C.** example of CBF signal spectrogram during ongoing activity at baseline and post-DCZ injection in a PV-hM4Di(Gi) mouse. **D**. Summary of average CBF at baseline and post-DCZ injection. **E**. Summary of average relative CBF fluctuation at baseline and post-DCZ injection. **F.** Average PSD of CBF signals at baseline and post-DCZ injection. **G.** Average BLPs of spontaneous CBF at baseline and post-DCZ injection. Significant differences are denoted by ** for p<0.01.

### DREADD activation

To activate hM4Di(Gi) or hM3Di(Gq) DREADD, we used DREADD agonist Deschloroclozapine (DCZ; HB85555, Hello Bio Inc, Princeton, NJ USA). 1 mg of DCZ was dissolved in 3.42 mL of 2% DMSO (prepared by diluting 100% DMSO with sterile saline). The solution was then aliquoted into ten 342.5 μL portions and stored at -20°C until use. For injections, aliquots were diluted with sterile saline to achieve a working concentration of 0.1 mg/mL. Mice were injected with 1 μg/kg DCZ dosage IP.

### Data Analysis

All data were analyzed using MATLAB (MathWorks, Natick, MA USA).

#### Cell region of interest (ROI)

Active cells expressing GCaMP were segmented from the average intensity 2P image using a custom-written function in MATLAB. The function segments cells based on correlation, identifying connected pixels based on temporal correlation with neighboring pixels. PV+ cells were identified based on mCherry labelling (Figure 1C), and the rest of the cells were considered PV-cells. The final ROIs were qualitatively examined to match the morphological features of neurons. The Ca^2+^ fluorescence signal of each cell body was measured by averaging all pixels within the ROI. Ca^2+^ change traces were corrected for neuropil contamination by regressing out the first two principal components of the neuropil Ca^2+^ trace.

#### Ca^2+^ event calculation

To infer Ca^2+^ events from the Ca^2+^ fluorescence traces, we applied OASIS (Online Active Set method to Infer Spikes) algorithm [27]. Fluorescence signal traces, normalized by mean fluorescence signal across time (ΔF/F), were deconvolved with the ‘deconvolveCa’ function, applying an autoregressive model of order 1 (’ar1’) with the ‘foopsi’ method, optimized with a sparsity penalty (’lambda’) set to 0.5 to balance the detection sensitivity for event amplitudes. Baseline fluorescence optimization (’optimize_b’) was enabled to correct for potential drifts, and the minimum spike threshold (’smin’) was set to -2.5.

#### LFP

Electrophysiology signals were obtained from EEG recordings. Raw EEG signals were filtered using a notch Fermi filter with cutoff frequencies of 58 Hz and 62 HZ and transition band of 1 Hz to remove 60 Hz power-line interference. The obtained signal was further filtered with a bandpass Fermi filter with cutoff frequencies of 1 Hz and 120 HZ and transition band of 1 Hz to obtain LFP activity. All LFP recordings were averaged across animals, normalized, and transformed into the frequency domain with mtspecgramc (Chronux toolbox), using tapers [2, 3] and a frequency range of 0 to 100 Hz. For each LFP signal, we computed power spectral density (PSD) using the fast Fourier transform (FFT). The frequency spectrum was obtained by taking the squared magnitude of the FFT of the signal, normalized by its length. For group comparisons, we averaged PSDs across animals and smoothed the log-transformed power values across frequency applying a moving average, focusing on the 0-100 Hz range. For each recording, the average BLP was calculated by dividing the power spectrum into six frequency bands (1-4 Hz, 5-8 Hz, 9-14 Hz, 15-29 Hz, 30-56 Hz, 64-100 Hz) using a bandpass Fermi filter with transition band of 1 Hz and computing the power percentage within each band relative to the total baseline power. For BLP time series, the resulting signals were squared, summed and low-pass filtered using a Fermi filter with cutoff frequency of 0.1 Hz and transition band of 0.1 Hz.

#### CBF

Time series of CBF changes were obtained from the LDF data. Raw CBF signals were low-pass filtered using a Fermi filter with cutoff frequency of 1 Hz and transition band of 1 Hz, down-sampled to 5 Hz and normalized by a mean of the baseline signal (CBF/CBF_0_). For each CBF signal, we computed the PSD in a similar manner as for LFP signal but focusing on the 0-1 Hz range. Average CBF was calculated as a mean of CBF/CBF_0_signal across time. Relative CBF fluctuation was calculated as standard deviation of CBF/CBF_0._ For each recording, the average BLP was calculated by dividing the power spectrum into three frequency bands (0.001-0.1 Hz, 0.1-1 Hz, and 0.001-1 Hz) using a bandpass Fermi filter with transition band of 1 Hz and computing the power percentage within each band relative to the total baseline power. For correlation analysis with LFP BLPs, CBF signals were low-pass filtered using a Fermi filter with cutoff frequency of 0.1 Hz and transition band of 0.1 Hz.

#### Diameter measurement

Vessel diameter was measured from 2P images as the full-width-at-half-maximum of the vessel cross-sectional intensity profile. Diameter changes were filtered using a Fermi filter with cutoff frequency of 1 Hz and transition band of 1 Hz.

#### Hemodynamic response function (HRF) fit

The HRF was defined by a gamma function with four parameters. The gamma fit parameters were determined using ‘lsqnonlin’ optimization routine in MATLAB to minimize the root mean squared error between the measured vessel diameter and neural input (GCaMP signal) convolved with the gamma function. The predicted vascular activity was obtained for each neuron by convolving GCaMP signal with HRF.

#### Apparent synchrony calculation

Apparent neuronal synchrony analysis was performed over overlapping 10-s time windows. For each window, we computed a correlation matrix for the spiking activity deconvolved with OASIS from Ca^2+^ fluorescence signals of all detected cells. To quantify synchrony, we calculated the mean of the upper triangular matrix values for each window. Only active cells (spiking probability > 1%) were included in this analysis. Synchrony peaks were identified by locating local maxima in the signal trace using a symmetric moving maximum filter with a 20-second window, selecting peaks sufficiently far from the signal edges, and extracting ±20-second segments around each peak for further analysis.

#### Statistical analysis

The Shapiro-Wilk test was applied to assess the normality of the data presented in this manuscript. We assessed significant differences in measurements between baseline and DCZ injection using a Mixed Design ANOVA (p<0.05 or p<0.01). For analyzing correlation between CBF and LFP, Pearson’s linear correlation coefficient was computed using MATLAB’s ‘corr’ function. For analyzing cross-correlation between vessel diameter changes and apparent network synchrony, cross-correlation coefficient was computed using MATLAB’s ‘xcorr’ function. We report average values ± standard deviation throughout the manuscript and figures, unless it is stated otherwise. No data were excluded when performing statistical analysis. No animals were excluded from analyses. The animal sample size for this study was determined based on a power analysis conducted using data from our previously published work. This analysis ensured that the sample size was sufficient to detect the expected effect size with adequate statistical power.

## RESULTS

### DREADD activation effectively modulates PV interneuron activity and alters network dynamics

We first confirmed that DREADD activation with DCZ injection effectively manipulated PV interneuron activity. Using 2P, we recorded ongoing neuronal population activity in awake, head-fixed mice expressing the Ca^2+^ sensor GCaMP (Figure 2A). An example of baseline activity from a single PV+ cell, represented by Ca^2+^ fluorescence changes, is shown in Figure 2B (black trace). Below, the raster plot illustrates Ca^2+^ activity in the PV+ cell population from the same animal, where each dot represents a Ca^2+^ event deconvolved from the corresponding Ca^2+^ fluorescence time series using OASIS algorithm [27].

DCZ injection in PV-hM4Di(Gi) mice effectively suppressed PV+ cell activity, as evidenced by a reduction in the number of Ca^2+^ fluorescence peaks and a lower number of Ca^2+^ events. This suppression was quantified as a significant decrease in average PV+ Ca^2+^ event frequency from 2.71 ± 0.13 events/min pre-DCZ to 1.85 ± 0.09 events/min post-DCZ (p = 2.810 × 10^-9^, Mixed Design ANOVA, 390 cells, 44 recordings, 9 animals, Figure 2D). However, the average Ca^2+^ event amplitude post-DCZ injection remained unchanged (0.21 ± 0.01 pre-DCZ vs. 0.21 ± 0.01 post-DCZ, p = 0.570, Mixed Design ANOVA, 390 cells, 44 recordings, 9 animals, Figure 2E).

Since PV interneurons regulate network activity through GABAergic inhibition of cortical excitatory neurons [16], we hypothesized that suppressing PV cell activity would increase overall network excitability. Consistent with this hypothesis, DCZ injection in PV-hM4Di(Gi) mice significantly increased PV-cell activity, as shown by elevated Ca^2+^ fluorescence peaks and an increased number of Ca^2+^ events. Average Ca^2+^ event frequency in PV-cells increased significantly from 1.78 ± 0.06 events/min pre-DCZ to 3.50 ± 0.10 events/min post-DCZ (p = 1.209 × 10^-66^, Mixed Design ANOVA, 806 cells, 44 recordings, 9 animals, Figure 2D). Additionally, the average Ca^2+^ event amplitude increased significantly from 0.21 ± 0.003 pre-DCZ to 0.25 ± 0.004 post-DCZ (p = 6.982 × 10^-18^, Mixed Design ANOVA, 806 cells, 44 recordings, 9 animals, Figure 2E).

Activating DREADD with DCZ in hM3Di(Gq) mice was also effective in enhancing PV cell activity, which led to a suppression of network excitability (Supplemental Figure 1). Control experiments confirmed that DCZ did not alter neuronal activity in the absence of DREADD expression, as DCZ injection in naïve mice had no effect on any neuronal sub-population (Supplemental Figure 2). These findings show that DCZ injection effectively activates DREADD, allowing precise manipulation of PV interneuron activity and network activity.

### Suppression of PV interneuron activity decreases high-gamma power and increases low-frequency power in LFP

Given the evidence that PV interneurons contribute to gamma activity, we hypothesized that suppressing PV neurons would reduce gamma power in LFP [17–18]. To test this, we recorded ongoing LFP activity in awake, head-fixed PV-hM4Di(Gi) mice using EEG (Figure 3A). An example of baseline LFP signals recorded in a single mouse before and after DCZ injection is shown in Figure 3B. The accompanying example spectrogram illustrates changes in power across frequencies over time (Figure 3C). Following DCZ injection, we observed a notable reduction in high-frequency power, quantified by calculating the BLPs (Supplemental Figure 3E, F). In the high-gamma range (64-100 Hz), the BLP significantly decreased from 0.89 ± 0.06% at baseline to 0.76 ± 0.06% post-DCZ injection (p = 1.834 × 10^-5^, Mixed Design ANOVA, 15 recordings, 7 animals, Figure 3E). However, in the gamma range (30-56 Hz), no significant change was observed (4.06 ± 0.2% at baseline vs. 4.12 ± 0.3% post-DCZ, p = 0.878, Mixed Design ANOVA, 15 recordings, 7 animals, Figure 3E).

Interestingly, suppressing PV cells also increased low-frequency BLPs (Supplemental Figure 3A-C). Specifically, delta (1-4 Hz) BLP increased significantly from 11.22 ± 0.41% at baseline to 23.96 ± 2.07% post-DCZ (p = 3.245 × 10^-4^, Mixed Design ANOVA, 15 recordings, 7 animals), theta (5-8 Hz) BLP increased from 17.14 ± 0.3% to 26.84 ± 1.49% (p = 3.662 × 10^-4^), and alpha (9-14 Hz) BLP increased from 9.11 ± 0.21% to 12.43 ± 0.90% (p = 0.023), whereas beta (15-29 Hz) BLP (6.72 ± 0.17% vs. 9.23 ± 0.96%, p = 0.082) did not show significant changes. We further verified that PV suppression also modulated sensory-evoked LFP activity by recording responses to whisker stimulation (50 ms pulses of air puffs delivered for 4 s every 40 s in 10 trials; Supplemental Figure 4A-B). These results are consistent with our findings from 2P Ca^2+^ imaging, where we observed increased excitatory cell (PV-) activity associated with delta and theta bands (Figure 2). Together, these findings demonstrate that suppressing PV interneurons decreases high-frequency power while increasing low-frequency power in LFPs, suggesting that PV suppression leads to network hyperexcitability.

### Suppression of PV interneuron activity alters spontaneous CBF

Previous studies on evoked activity have shown that PV interneurons contribute to CBF regulation [20–25]. Synchronized activation of PV cells in the S1 region induces a slow, delayed vasodilation in deeper cortical layers, where PV cells are predominantly located [21]. These slow vascular changes resemble the hemodynamic signals observed in resting-state studies in humans and animals. In this experiment, we investigated how suppressing PV interneurons affects spontaneous CBF fluctuations. We hypothesized that PV suppression would reduce the power of slow (low frequency) hemodynamic fluctuations. Using LDF, we recorded spontaneous CBF activity in awake, head-fixed mice (Figure 4A).

Examples of CBF signals recorded in a single PV-hM4Di(Gi) mouse during baseline conditions and post-DCZ injection and corresponding spectrograms are shown in Figure 4B-C. Suppression of PV cells with DCZ in PV-hM4Di(Gi) significantly increased the average normalized CBF signal from 0.99 ± 0.14×10^-4^ to 1.16 ± 0.17 (p = 0.002, Mixed Design ANOVA, 24 recordings, 9 animals, Figure 4D) and relative CBF fluctuation from 0.09 ± 0.02 to 0.13 ± 0.04 (p = 0.003, Mixed Design ANOVA, 24 recordings, 9 animals, Figure 4E). Surprisingly, PV suppression did not reduce the power of low-frequency CBF but increased the CBF power across all frequencies, as shown in PSD (Figure 4F). BLP analysis confirmed a significant increase in power within the low-frequency (0.001–0.1 Hz), high-frequency (0.1–1 Hz), and broad (0.001–1 Hz) bands (p = 0.004, p = 0.01, and p = 0.002, respectively; Mixed Design ANOVA, 24 recordings from 9 animals; Figure 4G). In contrast, activation of PV cells with DCZ in PV-hM3Di(Gq) mice significantly reduced the average CBF from 0.99 ± 0.67×10^-5^ to 0.93 ± 0.03 (p = 0.005, Mixed Design ANOVA, 4 recordings, 4 mice, Supplemental Figure 5D). However, the changes in the relative CBF fluctuation (p = 0.175, Mixed Design ANOVA, 4 recordings, 4 mice, Supplemental Figure 5E) and the power of CBF fluctuations (p = 0.139, p = 0.211, p = 0.226 for 0.001-0.1 Hz, 0.1-1 Hz, 0.001-1 Hz, respectively, Mixed Design ANOVA, 4 recordings, 4 mice, Supplemental Figure 5G) did not reach the significance level.

To determine whether the effects of PV suppression were local or global, we recorded CBF from the hemisphere opposite to the hM4Di(Gi) viral injection site. No changes in CBF were observed in the contralateral hemisphere (Supplemental Figure 6A). Additionally, in a control experiment with a naïve mouse (lacking DREADD), DCZ injection did not alter CBF (Supplemental Figure 6B). Overall, these results demonstrate that suppressing PV interneurons increases basal CBF, relative CBF fluctuation, and total CBF power.

### Suppression of PV interneuron activity increases basal arterial diameter and depth-dependent vascular dynamics

Because suppression of PV interneurons increased basal CBF in our LDF measurements, we next asked whether this effect was accompanied by dilation of individual arteries. Since PV interneurons are predominantly concentrated in layers 4 and 5 in the S1, we further hypothesized that PV modulation would produce the largest vascular changes in deeper layers [21]. To address this, we used 2P to measure depth-specific arterial diameters across cortical depths in awake PV-hM4Di(Gi) mice.

Figure 5A demonstrates an example of 2P intracortical sections with zoomed-in images of an artery in an awake mouse, which visibly increased in diameter following DCZ injection. This effect was further evident in the corresponding time profiles of arterial diameter changes (Figure 5B). In superficial layers (100-250 μm), suppression of PV cells with DCZ significantly increased the average basal arterial diameter from 13.00 ± 3.91 μm to 15.19 ± 4.44 μm (p=0.001, Mixed Design ANOVA, 10 recordings in 6 animals, Figure 5D top). In deeper layers (250-400 μm), we observed a similar trend in the average basal arterial diameter with the increase from 11.69 ± 4.06 μm to 13.92 ± 4.23 μm (p=0.005, Mixed Design ANOVA, 10 recordings in 6 animals, Figure 5D bottom). These results support our LDF measurements of increased CBF following PV suppression (Figure 4D). In contrast to LDF measurements showing decreased CBF with PV activation, DCZ injection in PV-hM3Di(Gq) mice did not significantly alter the average basal arterial diameter (16.91 ± 2.83 μm vs. 15.96 ± 2.58 μm; p = 0.114, Mixed Design ANOVA, 5 recordings from 5 animals, Supplemental Figure 7A, B, D).

**Figure 5.**
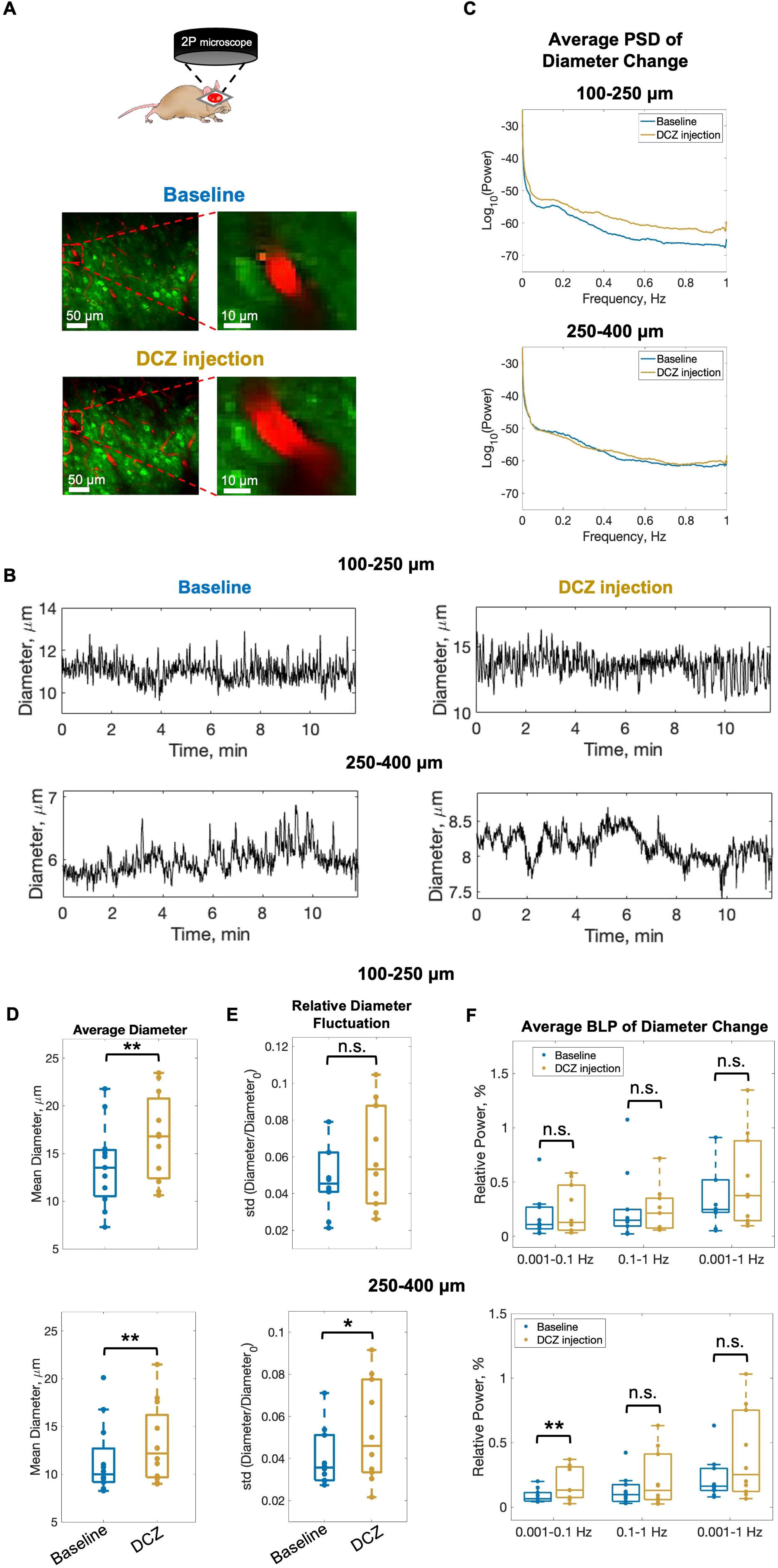
Suppression of PV neuron activity increases basal arterial diameter. **A.** Example 2P images of an artery located ∼250 μm below the cortical surface in a PV-hM4Di(Gi) mouse **B**. Example time profiles of artery diameter during ongoing activity at baseline and post-DCZ injection in superficial (top, 100-250 μm) and deeper (bottom, 250-400 μm) layers. **C**. Average PSD of artery diameter change at baseline and post-DCZ injection in superficial and deeper layers. Summary of **D.** average artery diameter, **E**. relative artery diameter fluctuation, and **F.** average BLPs of artery diameter change at baseline and post-DCZ injection in superficial (top) and deeper (bottom) layers. Significant differences are denoted by ** for p<0.01 and by * for p<0.05. Non-significant differences are denoted by n.s.

In contrast to superficial layers, where PV suppression did not significantly change relative arterial diameter fluctuation (0.05 ± 0.03 vs 0.07 ± 0.06, p=0.480, Mixed Design ANOVA, 10 recordings in 6 animals, Figure 5E top) and its power (p = 0.300, p = 0.706, p = 0.698 for 0.001-0.1 Hz, 0.1-1 Hz, and 0.001-1 Hz, respectively, Mixed Design ANOVA, 10 recordings in 6 animals, Figure 5F top), in deeper layers, it significantly increased the relative arterial diameter fluctuation from 0.04 ± 0.01 to 0.06 ± 0.02 (p=0.043, Mixed Design ANOVA, 10 recordings in 6 animals; Figure 5E bottom) and specifically the power of low-frequency (0.001-0.1 Hz) arterial diameter changes (p = 0.008, Mixed Design ANOVA, 10 recordings in 6 animals; Figure 5F bottom).The latter aligns with our LDF measurements, where PV suppression increased the power of CBF fluctuations. In summary, these findings show that suppressing PV interneurons increases the average basal arterial diameter across cortical depth and enhances low-frequency vascular fluctuations in deeper layers.

### Suppression of PV interneuron activity alters resting-state NVC

Previous studies in non-human primates and rodents have demonstrated a robust and consistent correlation between gamma frequency power and hemodynamic signals at rest [4, 8, 13–15]. Because PV suppression alters both network activity and CBF, we next examined whether it also impacts this strong gamma-CBF relationship. To address this, we simultaneously recorded ongoing resting-state LFP and CBF activity in awake, head-fixed PV-hM4Di(Gi) mice using EEG and LDF, respectively (Figure 6A).

**Figure 6.**
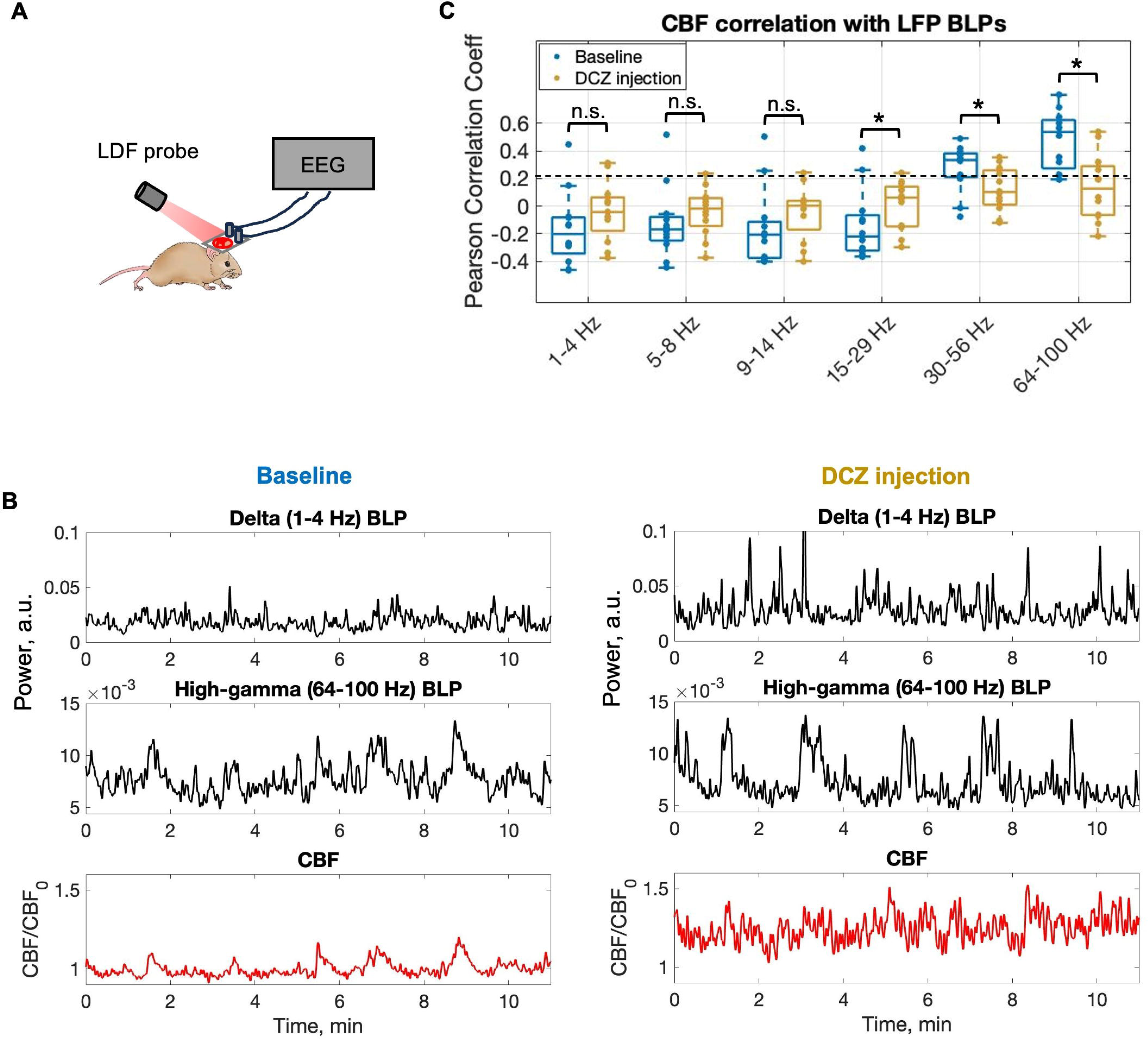
Suppression of PV neuron activity alters NVC. **A.** Setup for simultaneous recording of CBF and LFP with LDF and EEG, respectively. **B.** Example time profiles of delta band-limited power (1-4 Hz BLP), high-gamma band-limited power (64-100 Hz BLP), and cerebral blood flow (CBF) at baseline and post-DCZ injection in a single PV-hM4Di(Gi) mouse. **C**. CBF correlation with LFP BLPs in PV-hM4Di(Gi) mice at baseline and post-DCZ injection. Significant differences are denoted by * for p<0.05. Non-significant differences are denoted by n.s.

Examples of delta, high-gamma BLP, and CBF signals recorded in a single mouse under baseline conditions and post-DCZ injection are shown in Figure 6B. Our results align with previous studies, confirming that CBF changes are most strongly correlated with gamma (r = 0.169) and high-gamma (r = 0.276) LFP bands compared to delta (r = -0.158), theta (r = -0.159), alpha (r = -0.206), and beta (r = -0.110) bands (Figure 6C). Suppressing PV cells with DCZ significantly reduced the correlation between CBF and gamma BLP from 0.169 to 0.064 (p = 0.028, Mixed Design ANOVA, 12 recordings, 6 animals) and high-gamma BLP from 0.276 to 0.114 (p = 0.0002, Mixed Design ANOVA, 12 recordings, 6 animals). Interestingly, PV suppression significantly increased the CBF correlation with beta (15-29 Hz) BLP (p = 0.019, Mixed Design ANOVA, 12 recordings, 6 animals; Figure 6C). The CBF correlations with other frequency BLPs were not significantly affected (p = 0.087, p = 0.225, p = 0.158 for 1-4 Hz, 5-8 Hz, 9-14 Hz, respectively, Mixed Design ANOVA, 12 recordings, 6 animals; Figure 6C). Together, these findings show that suppressing PV interneurons disrupts the strong coupling between ongoing gamma activity and spontaneous low-frequency CBF fluctuations, while enhancing beta-band-coupling, suggesting a frequency-specific reorganization of resting-state NVC.

### Suppression of PV interneurons enhances neuronal synchrony while preserving coupling to arterial dynamics

Given that PV suppression produced hyper-excitability of excitatory neurons and enlarged basal arterial diameters, we investigated whether the resulting disinhibition alters the degree of network synchrony and its temporal coordination with spontaneous vascular dynamics. To test this, we quantified the synchrony of ongoing spiking activity, estimated from Ca²⁺ fluorescence traces via OASIS deconvolution, which we called “apparent neuronal synchrony”, and related it to spontaneous arterial diameter changes (Figure 7A). Apparent neuronal synchrony was calculated as the average pairwise correlation of spiking activity between active neurons within a 10 s moving window under baseline conditions and post-DCZ injection in PV-hM4Di(Gi) mice (Figure 7B, black trace).

**Figure 7.**
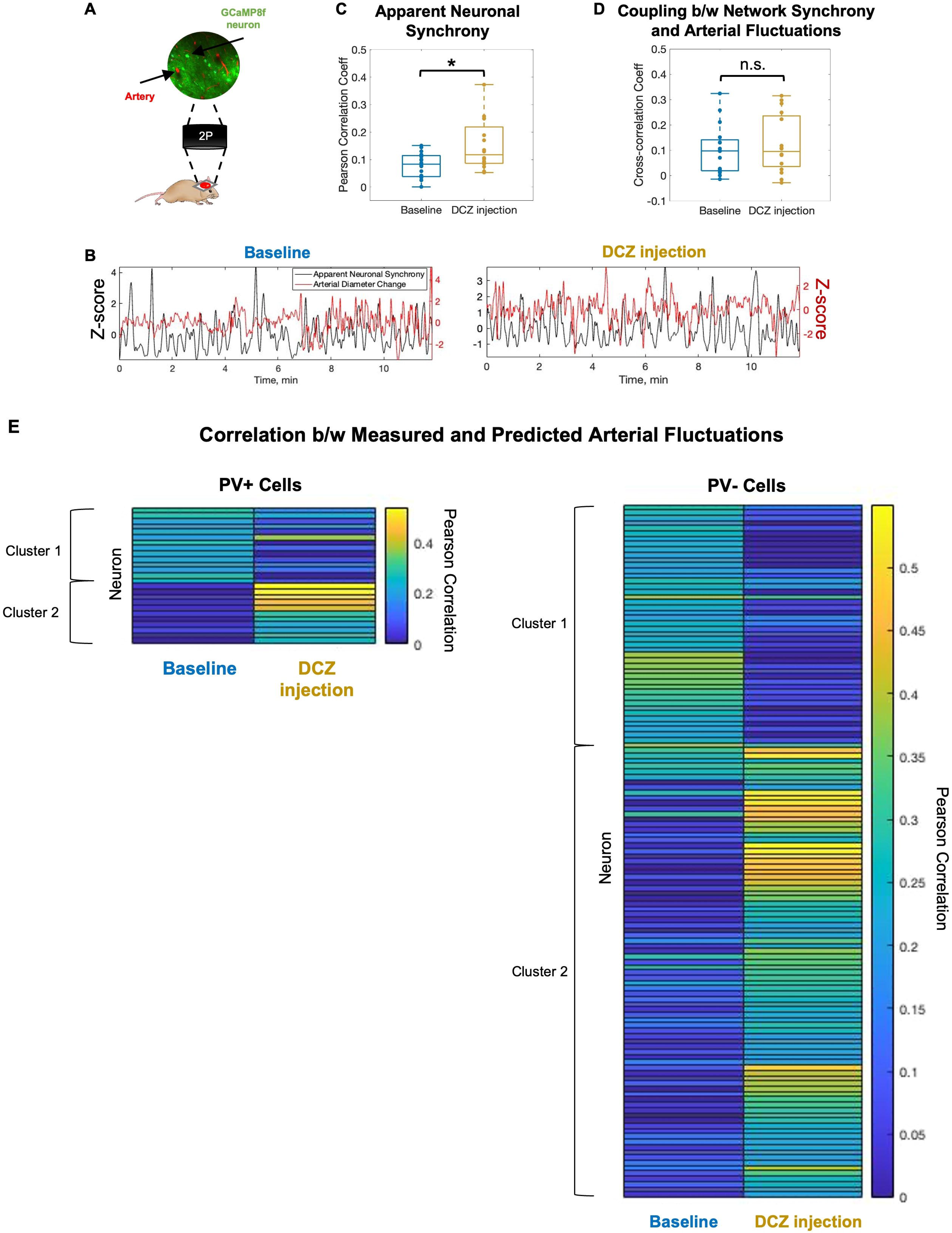
Suppression of PV neuron activity does not disrupt coupling between neuronal spiking synchrony and arterial fluctuations. **A.** Schematic of the 2P setup for simultaneous recording of neuronal Ca^2+^ signals (GCaMP fluorescence) and arterial diameter changes in the same field of view. **B**. Example time profiles of neuronal synchrony calculated as average spiking correlation between cells (black) and concurrent diameter changes of a nearby artery (red) in a PV-hM4Di(Gi) mouse at baseline and post-DCZ injection. **C**. Summary of apparent neuronal synchrony at baseline and post-DCZ injection. **D**. Summary of cross-correlation between apparent neuronal synchrony and spontaneous arterial diameter changes at baseline and post-DCZ injection. **E.** Heatmaps depicting Pearson correlation coefficients between measured arterial diameter changes and predictions based on PV+ or PV− cell activity, both at baseline and post-DCZ injection. Each row represents a single neuron, with rectangle colors indicating the magnitude of the correlation. For each neuronal subpopulation (PV+ and PV−), two clusters were identified: Cluster 1 comprises neurons with reduced correlation to arterial diameter changes after DCZ injection, while Cluster 2 includes neurons with increased correlation following DCZ injection.

PV suppression significantly increased average inter-neuronal correlation across animals (0.08 ± 0.01 to 0.15 ± 0.02; p = 0.013, paired t-test, 16 recordings from 16 animals; Figure 7C), suggesting an increase in apparent neuronal synchrony. We next examined the relationship between apparent neuronal synchrony and slow fluctuations in arterial diameter (Figure 7B, red trace). PV suppression did not significantly alter the mean cross-correlation between these signals (0.10 ± 0.02 baseline vs. 0.12 ± 0.03 post-DCZ; p = 0.570, paired t-test, 16 recordings from 16 animals; Figure 7D). These results suggest that despite disrupted gamma-CBF coupling upon PV suppression, the concurrent increase in beta-CBF coupling may have compensated, resulting in preserved coupling between broadband network synchrony and arterial dynamics.

### PV+ and PV− cells exhibit state-dependent contributions to vascular regulation

Since PV suppression did not disrupt the overall correlation between network synchrony and arterial fluctuations, we next asked whether specific neuronal subpopulations might compensate for altered network activity to maintain vascular regulation. To test this, we predicted arterial diameter changes from PV+ and PV-cell activity using a HRF fitted with a gamma function at the baseline and post-DCZ injection in awake PV-hM4Di(Gi) mice (Supplemental Figure 8). For each neuron, we computed the Pearson correlation between measured and predicted arterial diameter (Figure 7E). Neurons were selected from 20 recordings in 9 animals if the correlation exceeded 0.2 at either baseline or post-DCZ.

For PV+ cells, two clusters were identified based on baseline correlations: the first cluster (n=13 cells) had a higher average correlation (0.24 ± 0.03), while the second cluster (n=12 cells) had a lower average correlation (0.04 ± 0.06). Following DCZ injection, the first cluster’s correlation decreased to 0.08 ± 0.10 (p=0.005, n=13 cells), suggesting reduced contribution to vascular regulation, whereas the second cluster increased to 0.36 ± 0.12 (p=0.01, n=12 cells), suggesting increased contribution. Similarly, for PV-cells, baseline correlations revealed a high-correlation cluster (n=51 cells, 0.27 ± 0.05) and a low-correlation cluster (n=34 cells, 0.04 ± 0.07). Post-DCZ, the high-correlation cluster declined to 0.09 ± 0.12 (p=7.2×10^-13^, n=51 cells), while the low-correlation cluster increased to 0.32 ± 0.11 (p=1.662×^-14^, n=34 cells). Control analyses, in which 20 cells were randomly selected from each population, showed no clustering (Supplemental Figure 9), confirming that the observed patterns were not due to noise. These results indicate that both PV+ and PV− cells contribute to vascular regulation in a heterogeneous, state-dependent manner, differentiating normal versus hyperexcited network states. This may suggest that specific neuronal subpopulations may compensate when inhibitory input is reduced.

## DISCUSSION

Our findings reveal a complex role for PV interneurons in resting-state neurovascular dynamics, with distinct effects on network excitability, frequency-bounded activity, and cerebrovascular responses. By chemogenetically suppressing PV cells, we examined their contribution to neuronal activity, LFP power spectra, CBF, and vascular dynamics during ongoing activity. We confirmed the effective modulation of PV cells via DCZ activation of inhibitory or excitatory DREADDs using 2P Ca²⁺ imaging, which led to network hyperexcitability or suppression, respectively (Figure 2, Supplemental Figure 1). PV suppression reduced high-gamma power while increasing low-frequency LFP power (Figure 3, Supplemental Figure 3) and produced robust increase in both basal CBF and CBF fluctuations (Figure 4). Single-vessel 2P imaging supported these effects, showing enlarged basal arterial diameters and increased diameter fluctuations (Figure 5). Most importantly, PV suppression disrupted the strong coupling between gamma-band activity and spontaneous slow CBF fluctuations (Figure 6). At the cellular level, PV suppression significantly increased apparent neuronal synchrony without altering its coupling to spontaneous arterial responses, possibly due to compensation from specific neuronal subpopulations (Figure 7).

PV suppression caused a significant decrease in gamma-CBF correlation, suggesting that PV cell activity is essential for maintaining the canonical gamma-CBF relationship during resting state. This disruption occurred in the context of pronounced network hyperexcitability: excitatory cells showed elevated activity, and high-gamma power was reduced. Similar phenomena have been reported in Alzheimer’s disease conditions, where amyloid-β pathology disrupts excitation-inhibition balance, resulting in hyperactivity, hypersynchrony, and impaired oscillations [28–30]. Such functional disruptions often precede structural pathology, underscoring their role as a potential early biomarker of circuit dysfunction [30–31]. We were surprised not to observe a more significant reduction in gamma power following PV suppression. This outcome could be attributed to the elevated activity of excitatory cells, as increased spiking activity might elevate gamma power simply due to the additional energy generated.

The hemodynamics changes observed after PV suppression are likely secondary to elevated excitatory activity, as PV interneurons themselves are not major sources of vasoactive molecules. In contrast, excitatory neurons release vasoactive agents such as prostaglandin E_2_, which promote vasodilation [32]. Disinhibition likely enhances the release of these agents, sustaining the observed increases in basal CBF and arterial diameter. Conversely, PV activation, which reduces excitatory drive, decreased CBF without significantly altering basal arterial diameter, suggesting that reduced perfusion may arise from slower capillary flux rather than constriction of larger arteries. Together, these findings highlight the critical role of PV interneurons in maintaining both baseline cerebral perfusion and normal patterns of NVC.

Given the established role of gamma oscillations in local computation and feedforward signaling, the observed shift toward slower oscillations with elevated delta-alpha power likely reflects the emergence of synchronous population events under reduced inhibition [16–18]. Consistent with this, PV suppression increased apparent neuronal synchrony, as measured from Ca²⁺-inferred spiking activity. Disinhibition may promote the simultaneous recruitment of larger neuronal ensembles, increasing pairwise correlations across the network. While our synchrony metric, derived from deconvolved Ca²⁺ fluorescence signals, lacks the temporal precision of extracellular recordings, it provides reliable relative changes across conditions. The observed hypersynchrony aligns with previous findings in Alzheimer’s mouse model and epileptiform activity, where PV dysfunction or excessive excitation leads to paradoxical large-scale synchrony [28–30].

Interestingly, the coupling between apparent neuronal synchrony and spontaneous arterial diameter changes remained preserved despite hyperexcitability. The shift from gamma-CBF to beta-CBF coupling suggest that alternative oscillatory modes may compensate for the loss of gamma-mediated vascular regulation. Our HRF-based analysis further identified subsets of both PV+ and PV-cells that increased their contribution to arterial regulation after PV suppression, likely contributing to the preservation of synchrony-vascular coupling. These neurons may maintain perfusion by releasing vasoactive molecules or engaging alternative signaling pathways, effectively compensating for reduced inhibitory control. This adaptability suggests the resilience of cortical circuits in preserving vascular homeostasis under altered excitation-inhibition balance. Future studies should characterize these compensatory neuronal subtypes in greater detail, including their molecular identity, synaptic connectivity, and capacity to regulate vascular tone under physiological and pathological conditions. Understanding how such adaptative mechanisms operate could inform therapeutic strategies aimed at restoring healthy NVC in disorders associated with PV dysfunction.

In summary, our results provide mechanistic insight into the cellular basis of spontaneous low-frequency hemodynamic fluctuations that underpin resting-state functional connectivity. We demonstrate that PV interneurons are critical for maintaining the canonical gamma-CBF coupling that likely supports the temporal coherence of vascular signals across distant but functionally connected brain regions. Disruption of PV activity not only alters local oscillatory-vascular relationships but also reorganizes the frequency content of NVC, shifting toward beta–CBF correlations. By identifying PV interneurons as key modulators of the resting-state vascular signature, our study advances the interpretability of hemodynamic-based imaging measures, helping to detect changes in functional connectivity arising from altered neuronal dynamics. This mechanistic understanding is essential for accurately linking fMRI and other hemodynamic imaging readouts to the underlying neural processes in both health and disease.

## ACKNOWLEDGEMENTS

We thank Dr. Ping Wang and Xiaoling Yang for their technical support. We also thank the University of North Carolina Viral Vector Core for generously providing and facilitating access to the chemogenetic constructs used in this work. We thank the GENIE project (Janelia) for making calcium sensors available for use.

## AUTHOR CONTRIBUTIONS

Conceptualization, A.R. and A.L.V.; Methodology, A.R., M. F., and A.L.V.; Validation, A.R.; Formal Analysis, A.R. and A.L.V.; Investigation, A.R.; Resources, M. F. and A.L.V.; Writing – Original Draft, A.R.; Writing - Review & Editing, A.R., M. F., and A.L.V.; Visualization: A.R.; Supervision, A.L.V.; Project Administration, A.R. and A.L.V.; Funding Acquisition, A.L.V.

## DATA AVAILABILITY

All data that support the findings of this study are available from the lead contact upon request. Processed calcium imaging, optogenetic, and chemogenetic mouse data have been deposited at GitHub and are publicly available as of the date of publication.

## SUPPLEMENTAL INFORMATION

Document S1. Figures S1-S9.

## REFERENCES

[1] Biswal B, Yetkin FZ, Haughton VM, Hyde JS. Functional connectivity in the motor cortex of resting human brain using echo-planar MRI. Magn Reson Med. 1995 Oct;34(4):537–41.

[2] Fox MD, Raichle ME. Spontaneous fluctuations in brain activity observed with functional magnetic resonance imaging. Nat Rev Neurosci. 2007 Sep;8(9):700–11.

[3] Drew PJ, Mateo C, Turner KL, Yu X, Kleinfeld D. Ultra-slow Oscillations in fMRI and Resting-State Connectivity: Neuronal and Vascular Contributions and Technical Confounds. Neuron. 2020 Sep 9;107(5):782–804.

[4] Mateo C, Knutsen PM, Tsai PS, Shih AY, Kleinfeld D. Entrainment of Arteriole Vasomotor Fluctuations by Neural Activity Is a Basis of Blood-Oxygenation-Level-Dependent “Resting-State” Connectivity. Neuron. 2017 Nov 15;96(4):936–948.e3.

[5] De Luca M, Beckmann CF, De Stefano N, Matthews PM, Smith SM. fMRI resting state networks define distinct modes of long-distance interactions in the human brain. Neuroimage. 2006 Feb 15;29(4):1359–67.

[6] Sepulcre J, Liu H, Talukdar T, Martincorena I, Yeo BTT, Buckner RL. The organization of local and distant functional connectivity in the human brain. PLoS Comput Biol. 2010 Jun 10;6(6):e1000808.

[7] Ma Y, Shaik MA, Kozberg MG, Kim SH, Portes JP, Timerman D, et al. Resting-state hemodynamics are spatiotemporally coupled to synchronized and symmetric neural activity in excitatory neurons. Proc Natl Acad Sci USA. 2016 Dec 27;113(52):E8463–71.

[8] Murphy MC, Chan KC, Kim S-G, Vazquez AL. Macroscale variation in resting-state neuronal activity and connectivity assessed by simultaneous calcium imaging, hemodynamic imaging and electrophysiology. Neuroimage. 2018 Apr 1;169:352–62.

[9] Yeo BTT, Krienen FM, Sepulcre J, Sabuncu MR, Lashkari D, Hollinshead M, et al. The organization of the human cerebral cortex estimated by intrinsic functional connectivity. J Neurophysiol. 2011 Sep;106(3):1125–65.

[10] Greicius MD, Krasnow B, Reiss AL, Menon V. Functional connectivity in the resting brain: a network analysis of the default mode hypothesis. Proc Natl Acad Sci USA. 2003 Jan 7;100(1):253–8.

[11] Logothetis NK, Pauls J, Augath M, Trinath T, Oeltermann A. Neurophysiological investigation of the basis of the fMRI signal. Nature. 2001 Jul 12;412(6843):150–7.

[12] Leopold DA, Murayama Y, Logothetis NK. Very slow activity fluctuations in monkey visual cortex: implications for functional brain imaging. Cereb Cortex. 2003 Apr;13(4):422–33.

[13] Shmuel A, Leopold DA. Neuronal correlates of spontaneous fluctuations in fMRI signals in monkey visual cortex: Implications for functional connectivity at rest. Hum Brain Mapp. 2008 Jul;29(7):751–61.

[14] Schölvinck ML, Maier A, Ye FQ, Duyn JH, Leopold DA. Neural basis of global resting-state fMRI activity. Proc Natl Acad Sci USA. 2010 Jun 1;107(22):10238–43.

[15] Winder AT, Echagarruga C, Zhang Q, Drew PJ. Weak correlations between hemodynamic signals and ongoing neural activity during the resting state. Nat Neurosci. 2017 Dec;20(12):1761–9.

[16] Bartos M, Vida I, Jonas P. Synaptic mechanisms of synchronized gamma oscillations in inhibitory interneuron networks. Nat Rev Neurosci. 2007 Jan;8(1):45–56.

[17] Cardin JA, Carlén M, Meletis K, Knoblich U, Zhang F, Deisseroth K, et al. Driving fast-spiking cells induces gamma rhythm and controls sensory responses. Nature. 2009 Jun 4;459(7247):663–7.

[18] Sohal VS, Zhang F, Yizhar O, Deisseroth K. Parvalbumin neurons and gamma rhythms enhance cortical circuit performance. Nature. 2009 Jun 4;459(7247):698–702.

[19] Vazquez AL, Fukuda M, Kim S-G. Inhibitory neuron activity contributions to hemodynamic responses and metabolic load examined using an inhibitory optogenetic mouse model. Cereb Cortex. 2018 Nov 1;28(11):4105–19.

[20] Krawchuk MB, Ruff CF, Yang X, Ross SE, Vazquez AL. Optogenetic assessment of VIP, PV, SOM and NOS inhibitory neuron activity and cerebral blood flow regulation in mouse somato-sensory cortex. J Cereb Blood Flow Metab. 2020 Jul;40(7):1427–40.

[21] Rakymzhan A, Fukuda M, Kozai TDY, Vazquez AL. Parvalbumin interneuron activity induces slow cerebrovascular fluctuations in awake mice. BioRxiv. 2024 Jun 16. DOI: 10.1101/2024.06.15.599179.

[22] Dahlqvist MK, Thomsen KJ, Postnov DD, Lauritzen MJ. Modification of oxygen consumption and blood flow in mouse somatosensory cortex by cell-type-specific neuronal activity. J. Cereb. Blood Flow Metab. 2020 Oct;40(10):2010–25. doi: 10.1177/0271678X19882787.

[23] Lee J, Stile CL, Bice AR, Rosenthal ZP, Yan P, Snyder AZ, Lee J-M, Bauer AQ. Opposed hemodynamic responses following increased excitation and parvalbumin-based inhibition. J Cereb Blood Flow Metab 2021;41:841–56.

[24] Krogsgaard A, Sperling L, Dahlqvist M, Thomsen K, Vydmantaite G, Li F, et al. PV interneurons evoke astrocytic Ca2+ responses in awake mice, which contributes to neurovascular coupling. Glia. 2023 Mar 30; 71,8 (2023): 1830–1846.

[25] Vo TT, Im GH, Han K, Suh M, Drew PJ, Kim SG. Parvalbumin interneuron activity drives fast inhibition-induced vasoconstriction followed by slow substance P-mediated vasodilation. Proc. Natl. Acad. Sci. U.S.A. 2023 Apr;120(18):e2220777120.

[26] Masamoto K, Tomita Y, Toriumi H, Aoki I, Unekawa M, Takuwa H, Itoh Y, Suzuki N, Kanno I.Repeated longitudinal in vivo imaging of neuro-glio-vascular unit at the peripheral boundary of ischemia in mouse cerebral cortex. Neuroscience. 2012 Jun 14;212:190–200.

[27] Friedrich J, Zhou P, Paninski L. Fast online deconvolution of calcium imaging data. PLoS computational biology. 2017 Mar 14;13(3):e1005423.

[28] Verret L, Mann EO, Hang GB, Barth AMI, Cobos I, Ho K, et al. Inhibitory interneuron deficit links altered network activity and cognitive dysfunction in Alzheimer model. Cell. 2012 Apr 27;149(3):708–21.

[29] Palop JJ, Mucke L. Network abnormalities and interneuron dysfunction in Alzheimer disease. Nature Reviews Neuroscience. 2016 Dec;17(12):777–92.

[30] Maestú F, de Haan W, Busche MA, DeFelipe J. Neuronal excitation/inhibition imbalance: core element of a translational perspective on Alzheimer pathophysiology. Ageing Res Rev. 2021 Aug;69:101372.

[31] Leitch B. Parvalbumin interneuron dysfunction in neurological disorders: focus on epilepsy and alzheimer’s disease. Int J Mol Sci. 2024 May 19;25(10).

[32] Lecrux C, Toussay X, Kocharyan A, Fernandes P, Neupane S, Lévesque M, et al. Pyramidal neurons are “neurogenic hubs” in the neurovascular coupling response to whisker stimulation. J Neurosci. 2011 Jul 6;31(27):9836–47.

