## Supplemental Figures for "Parvalbumin interneurons mediate spontaneous hemodynamic fluctuations"

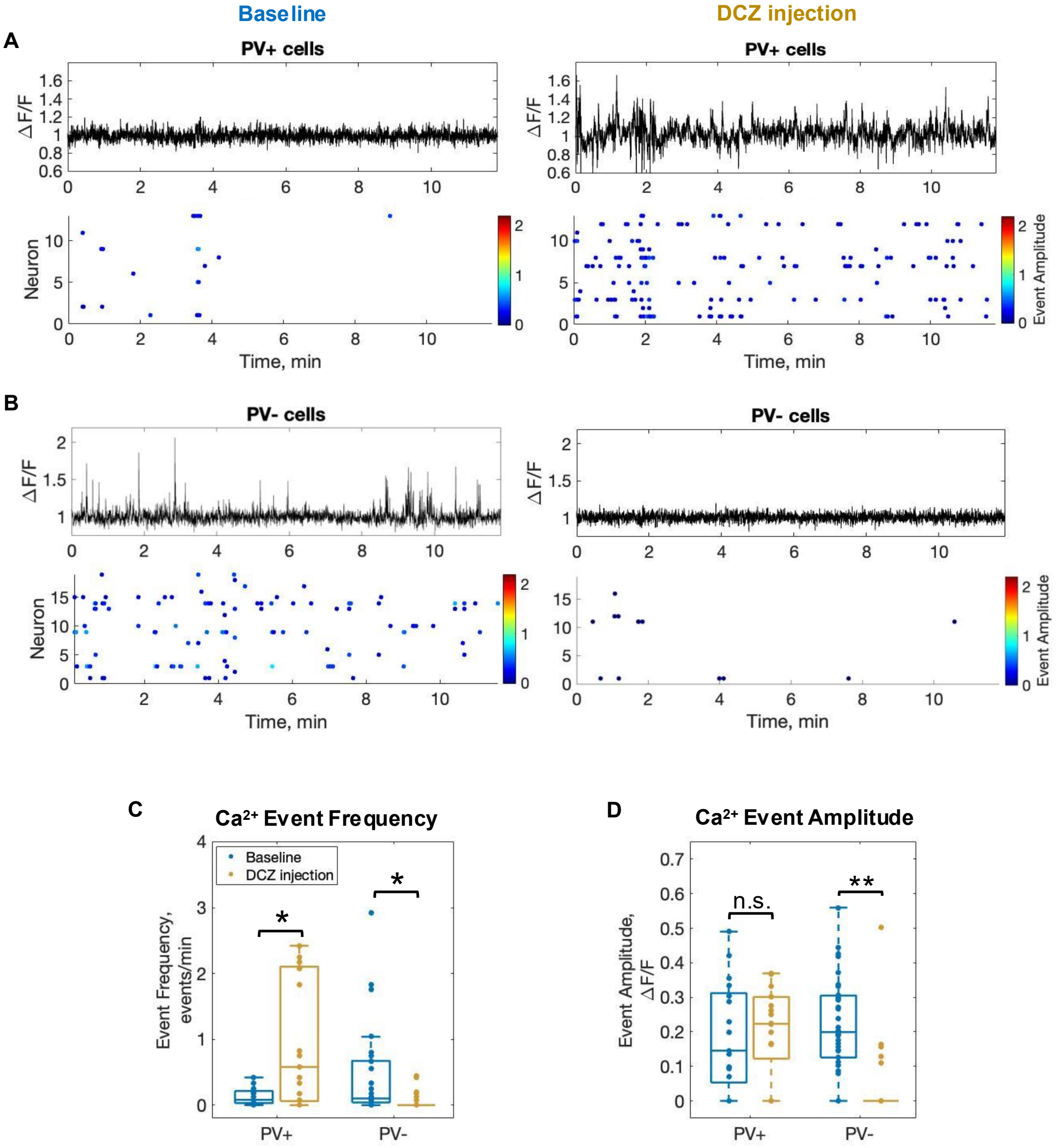


**Supplemental Figure 1. Activation of hM3Di(Gq) DREADD with DCZ effectively enhances PV neuron activity and suppresses network activity. A.** Time profiles of $Ca^{2+}$ fluorescence changes (black traces) in an individual PV+ cell in a PV-hM3Di(Gq) mouse during ongoing activity at baseline and post-DCZ injection. Below are the raster plots of $Ca^{2+}$ activity in PV+ cell population from a single animal at the baseline and post-DCZ injection. Each dot represents a $Ca^{2+}$ event and each raw represents an individual neuron. The dot color encodes an event amplitude. **B**. Similar analysis as in A but for PV- cells. **C**. Summary of $Ca^{2+}$ event frequency and **D**. $Ca^{2+}$ event amplitude in 34 PV+ cells (frequency: p=0.016, Mixed Design ANOVA; amplitude: p=0.73, Mixed Design ANOVA) and 68 PV- cells (frequency: p=0.015, Mixed Design ANOVA; amplitude: $p=7.95*10^{-5}$, Mixed Design ANOVA) identified from 3 recordings in 3 animals. Significant differences are denoted by * for p<0.05 or ** for p<0.01. Non-significant differences are denoted by n.s.

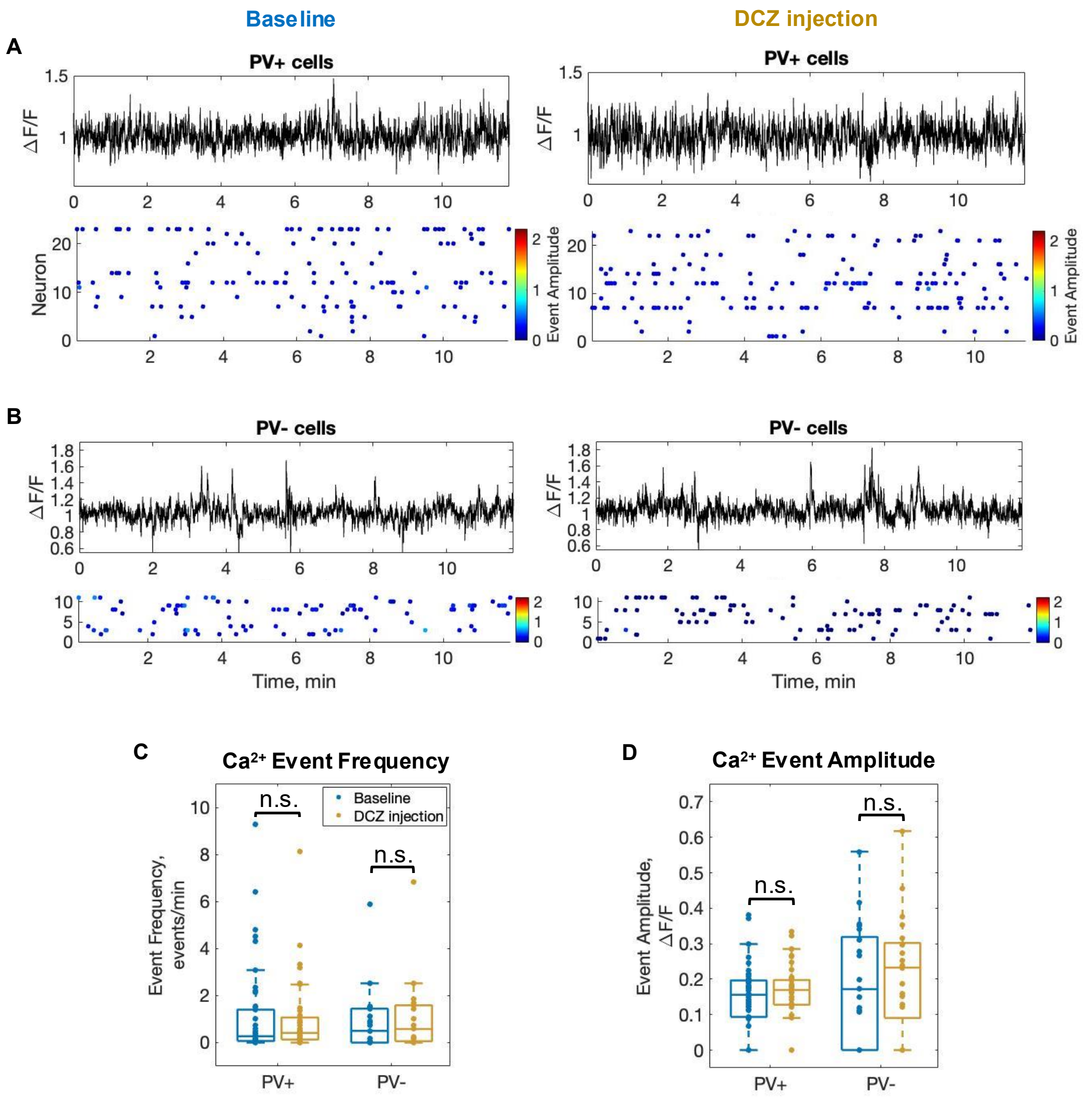


**Supplemental Figure 2. DCZ injection does not affect neuronal activity in naïve mice. A.** Time profiles of $Ca^{2+}$ fluorescence changes (black traces) in an individual PV+ cell in a PV-mCherry mouse (from a control group) during ongoing activity at baseline and post-DCZ injection. Below are the raster plots of $Ca^{2+}$ activity in PV+ cell population from a single animal at baseline and post-DCZ injection. Each dot represents a $Ca^{2+}$ event and each raw represents an individual neuron. The dot color encodes an event amplitude. **B**. Similar analysis as in A but for PV- cells. **C**. Summary of $Ca^{2+}$ event frequency and **D**. $Ca^{2+}$ event amplitude in 92 PV+ cells and 42 PV- cells identified from 3 recordings in 3 animals (control group). No significant differences between baseline and post-injection are detected for event frequency or event amplitude in any neuronal sub-population. Non-significant differences are denoted by n.s.

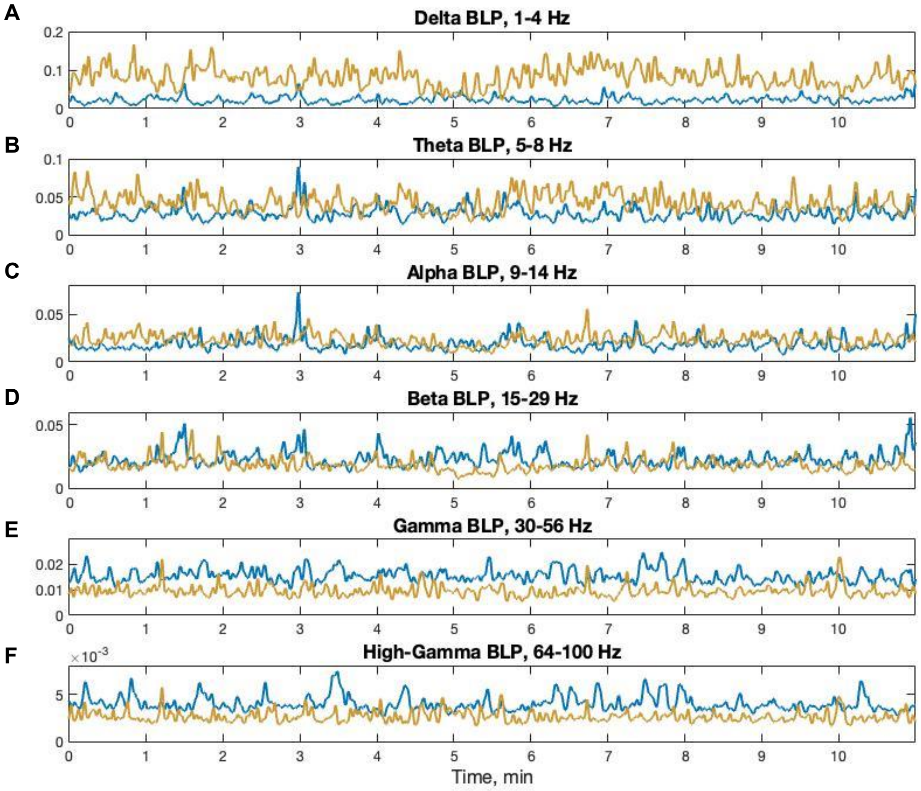


**Supplemental Figure 3. Effect of suppressing PV neuron activity on band-limited LFP power.** Example time courses of BLPs at baseline (blue) and post-DCZ injection (yellow) in a PV-hM4Di(Gi) mouse.

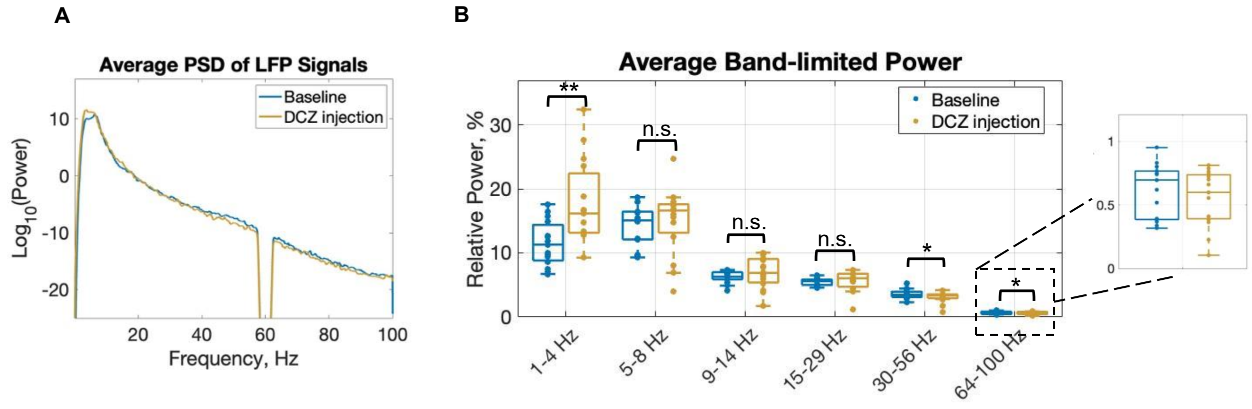


**Supplemental Figure 4. Effect of suppressing PV neuron activity on LFP power during sensory stimulation. A.** Average PSD of LFP signals in PV-hM4Di(Gi) mice (15 recordings in 7 animals) recorded during sensory stimulation at baseline and post-DCZ injection. **B**. Average LFP BLPs in PV-hM4Di(Gi) mice at baseline and post-DCZ injection (15 recordings in 7 animals) recorded during sensory stimulation. 1-4 Hz, p=0.002; 4-8 Hz, p=0.552; 8-14 Hz, p=0.302; 14-30 Hz, p=0.718; 30-56 Hz, p=0.017; 64-100 Hz, p=0.030, Mixed Design ANOVA. Significant differences are denoted by * for p<0.05 or ** for p<0.01. Non-significant differences are denoted by n.s.

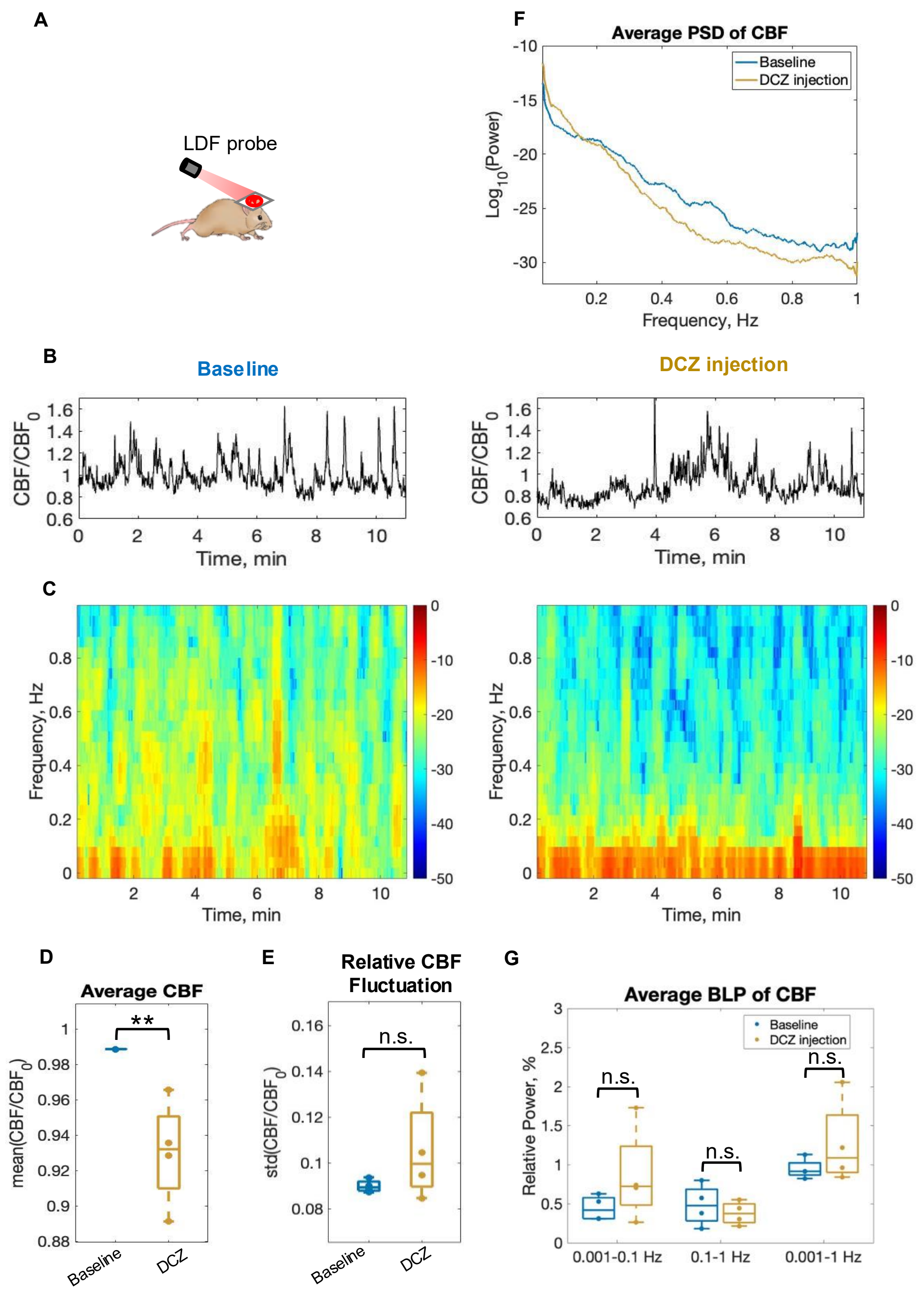


**Supplemental Figure 5. Activation of PV neuron activity decreases spontaneous CBF. A.** LDF recording setup for CBF measurements, where an LDF probe was placed on top of a cranial window. **B.** Example of CBF time profiles and. **C.** example of CBF signal spectrogram during ongoing activity at baseline and post-DCZ injection in a PV-hM3Di(Gq) mouse. **D.** Summary of average CBF at baseline and post-DCZ injection. **E.** Summary of average relative CBF fluctuation at baseline and post-DCZ injection. **F.** Average PSD of CBF signals at baseline and post-DCZ injection. **G.** Average BLPs of spontaneous CBF at baseline and post-DCZ injection. 0.001-0.1 Hz, $p=0.139$; 0.1-1 Hz, $p=0.211$; 0.001-1 Hz: $p=0.226$. Significant differences are denoted by ** for $p<0.01$. Non-significant differences are denoted by n.s.

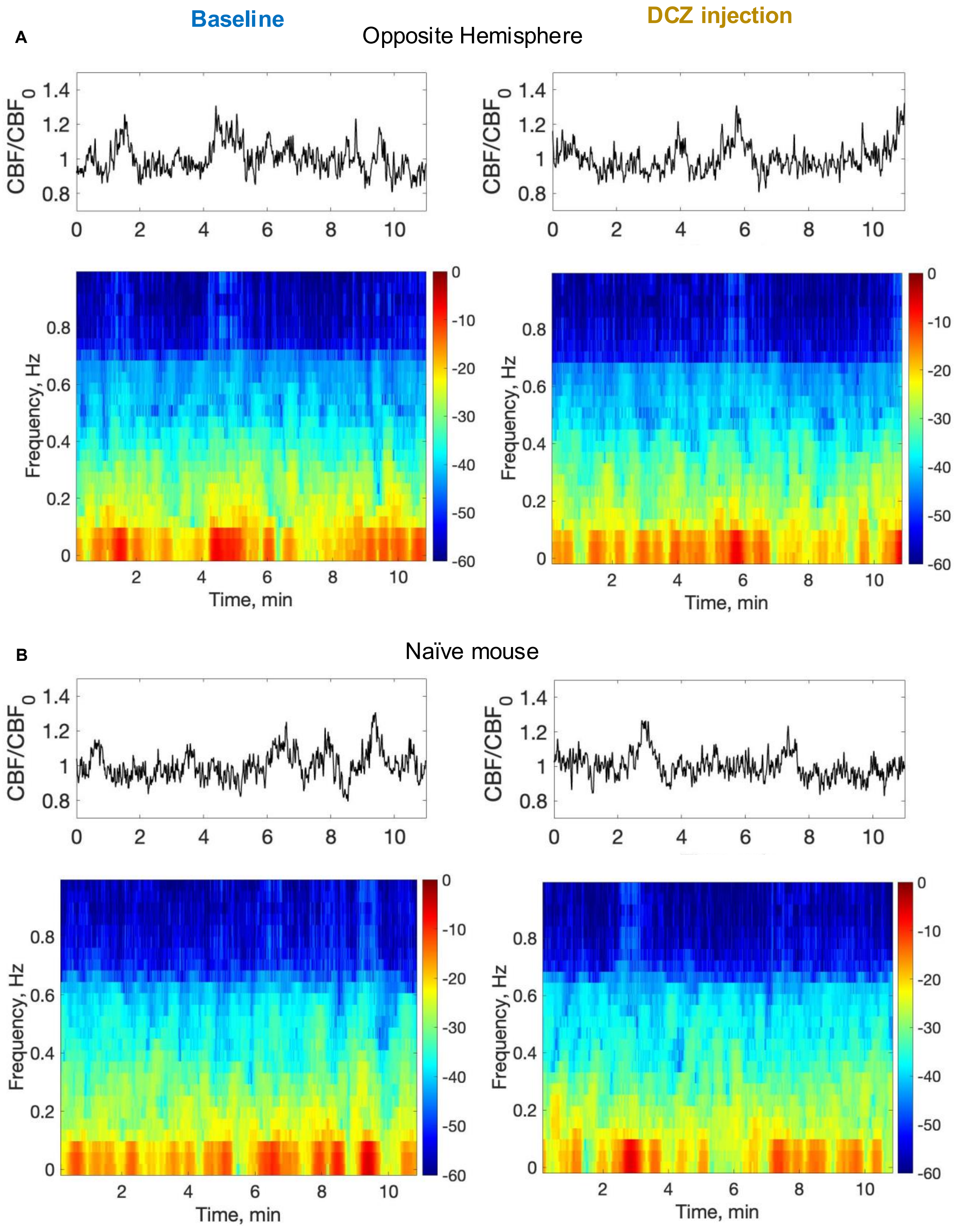


**Supplemental Figure 6. DCZ injection does not change CBF in the control group. A.** Example of spontaneous CBF time profiles and corresponding spectrograms in the opposite hemisphere of a PV-hM4Di(Gi) mouse and **B**. in a PV-mCherry (naïve) mouse at the baseline and post-DCZ injection.

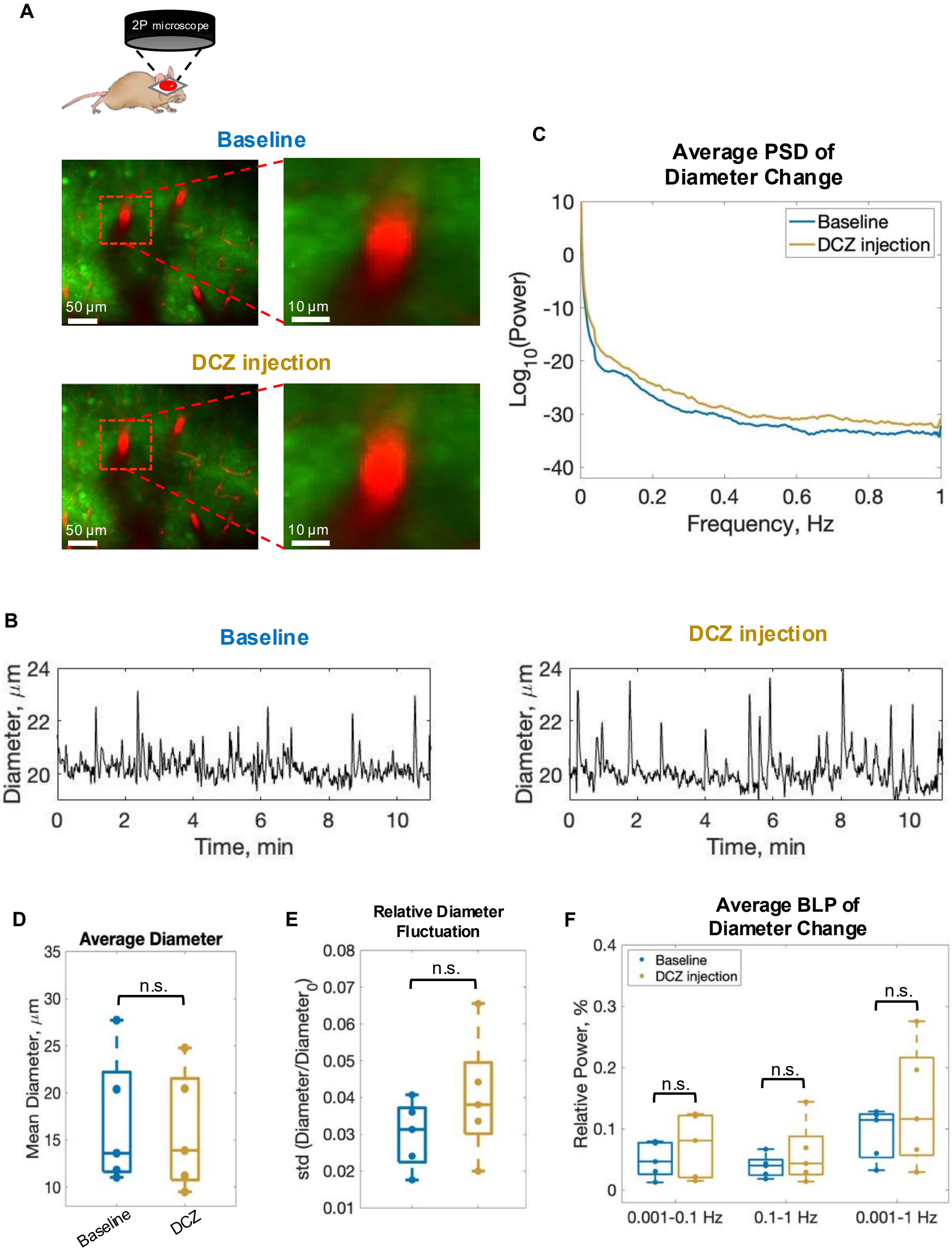


**Supplemental Figure 7. Activation of PV neuron activity does not change basal arterial diameter. A.** Example 2P images of an artery located ~250 μm below the cortical surface in a PV-hM3Di(Gq) mouse **B**. Example time profiles of artery diameter during ongoing activity at baseline and post-DCZ injection. **C**. Average power spectral density of artery diameter change at baseline and post-DCZ injection. **D.** Summary of average artery diameter and. **E**. relative artery diameter fluctuation at baseline and post-DCZ injection. **F.** Average band-limited powers of artery diameter change at baseline and post-DCZ injection. Non-significant differences are denoted by n.s.

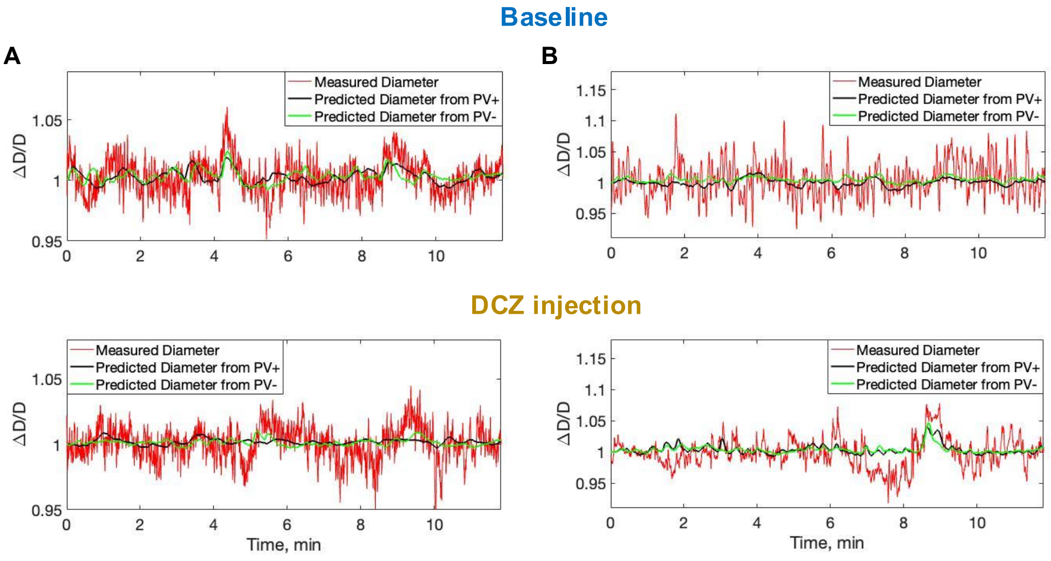


**Supplemental Figure 8. Measured vs. predicted arterial diameter changes. A.** Example time profiles showing measured arterial diameter changes (red), predicted diameter changes based on activity from a single PV+ cell (black), and predicted diameter changes based on activity from a single PV- cell (green) for a single vessel in a PV-hM4Di(Gi) mouse at baseline and post-DCZ injection. In this example, the correlation between measured and predicted (from both PV+ and PV-) diameters decreased post-DCZ injection. **B**. Similar example as in B, where the correlation between measured and predicted (from both PV+ and PV-) diameters increased post-DCZ injection.

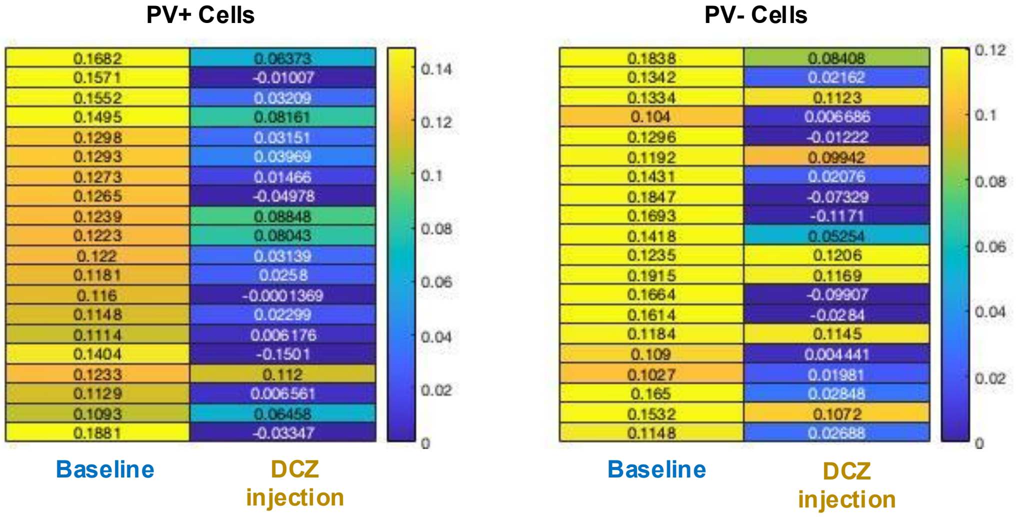


**Supplemental Figure 9. The observed cell clustering was not the effect of noise.** Heatmaps depicting Pearson correlation coefficients between arterial diameter changes and predictions based on activity of randomly selected 20 PV+ or PV− cells, both at baseline and post-DCZ injection. Each row represents a single neuron, with rectangle colors indicating the magnitude of the correlation. No definitive cluster were identified for either neuronal subpopulation (PV+ and PV−).
